# Nodule-specific Cu^+^-chaperone NCC1 is required for symbiotic nitrogen fixation in *Medicago truncatula* root nodules

**DOI:** 10.1101/2023.03.05.531139

**Authors:** Cristina Navarro-Gómez, Javier León-Mediavilla, Hendrik Küpper, Mario Rodríguez-Simón, Alba Paganelli-López, Jiangqi Wen, Stefan Burén, Kirankumar S. Mysore, Syed Nadeem Hussain Bokhari, Juan Imperial, Viviana Escudero, Manuel González-Guerrero

## Abstract

Cu^+^-chaperones are a diverse group of proteins that allocate Cu^+^ ions to specific copper-proteins, creating different copper pools targeted to specific physiological processes. Symbiotic nitrogen fixation carried out in legume root nodules indirectly requires relatively large amounts of copper e.g. for energy delivery via respiration, for which targeted copper deliver systems would be required. MtNCC1 is a nodule-specific Cu^+^-chaperone encoded in the *Medicago truncatula* genome, with a N-terminus Atx1-like domain that can bind Cu^+^ with picomolar affinities. This gene is expressed primarily from the late infection zone to the early fixation zone, and is located in the cytosol, associated to plasma and symbiosome membranes, and within nuclei. Consistent with its key role in nitrogen fixation, *ncc1* mutants have a severe reduction of nitrogenase activity, and a 50% reduction in copper-dependent cytochrome *c* oxidase activity. A subset of the copper-proteome is also affected in the mutant nodules. Many of these proteins can be pulled-down when using a Cu^+^-loaded N-terminal MtNCC1 moiety as a bait, indicating a role in nodule copper homeostasis and in copper-dependent physiological processes. Overall, these data suggest a pleiotropic role of MtNCC1 in copper delivery for symbiotic nitrogen fixation.

## INTRODUCTION

Copper is an essential nutrient for plants (Marschner and Marschner, 2011). It is involved in key physiological processes such as photosynthesis, respiration, ethylene signalling, or free radical control, among many others (Andresen et al, 2018). This versatile use of copper is largely based on its ability to transition between two redox states (Cu^+^ and Cu^2+^) in physiological conditions (Burkhead et al., 2009). However, that same property makes copper a toxic reagent at slightly higher concentration (Küpper and Andresen, 2016). One suggested mechanism of toxicity is the non-enzymatic catalysis of Fenton-style reactions producing damaging free radicals (Goldstein et al., 1993). Furthermore, excess copper can displace other essential transition metals from the active site of metalloproteins (Küpper et al., 1996; Küpper et al., 2002; Macomber and Imlay, 2009). As a result, and to prevent copper damage to the cell structures, the “free”, hydrated, concentrations of copper in the cytosol are maintained at extremely low levels, less than one ion per cell (Changella, 2003). This is achieved by the coordinated action of small, copper-binding molecules, as well as proteins known as Cu^+^-chaperones (Robinson and Winge, 2010; Flis et al., 2016).

The role of Cu^+^-chaperones is to deliver copper to different copper-proteins, protecting the cell from copper toxicity (Lin et al., 1997; Wong et al., 2000). Cu^+^-chaperones bind Cu^+^ with picomolar (pM)-femtomolar (fM) affinity (Rae et al., 1999; Palumaa et al., 2004), which prevents unspecific copper release prior to docking with a compatible protein. This means that metalation is not simply the result of relative copper-binding affinities, but also of the specific protein-protein interaction between donor and acceptor proteins. Consequently, different copper pools are created based on the ability of each protein to interact and exchange copper with one or another Cu^+^-chaperone. *Saccharomyces cerevisiae* has at least three of these pools: the secretory pathway that receives copper from Atx1 (Lin et al., 1997); the mitochondria that obtain copper from COX17 (Palumaa et al., 2004); and the Cu, Zn superoxide dismutase that gets it from CCS (Rae et al., 1999). Multicellular organisms are expected to have a larger number of such proteins, which are related in structure but serve a diversity of functions. Plants have at least a CCS orthologue (Chu et al., 2005), two Cu^+^-chaperones driving copper to mitochondria (Attallah et al., 2011), another in the stroma (Blaby-Haas et al., 2014), and three ATX1 orthologues with the characteristic Cu^+^-binding CXXC motive: ATX1 (Shin et al., 2012), CCP (Chai et al., 2020) and CCH (Mira et al., 2001), the later with a C-terminal domain of unknown function. This multiplicity of Cu^+^-chaperones hints not only at the existence of different copper pools, but also at their specialized roles in plant physiology. For instance, ATX1 seems to be involved in buffering through copper deficiency or excess (Shin et al., 2012), CCH has been proposed to be involved in recovering copper during leave senescence (Mira et al., 2001), and CCP plays a role in plant immunity (Chai et al., 2020). Therefore, it should be expected that other, new Cu^+^-chaperones are associated with other copper-dependent processes.

While leaves are the main copper sink in most plants during vegetative growth, legumes have a second, major copper sink in their root nodules (Johnston et al., 2001; Senovilla et al., 2018). After the exchange of specific signals between legumes and a group of bacteria known as rhizobia, cells in the root cortex, pericycle and endodermis proliferate to produce nodules (Downie, 2014; Xiao et al., 2014). Concomitant to nodule development, rhizobia penetrate and colonize nodule cells. Surrounded by the plant-derived symbiosome membrane, rhizobia differentiate into bacteroids that synthesize nitrogenase to fix nitrogen. This developmental process can be easily followed in indeterminate-type nodules, such as those in pea or in *Medicago truncatula*. Indeterminate nodules maintain their apical meristem over time, leading to a spatial-temporal gradient. As a result, four developmental zones can be observed in these nodules: the apical meristem (zone I), the infection-differentiation zone (zone II), an interzone where oxygen levels drop to prevent nitrogenase inhibition, so that nitrogen fixation can occur in the fixation zone of the nodule (zone III), and a senescent zone (zone IV) in older nodules (Vasse et al., 1990). In exchange for this fixed nitrogen, the host plant provides photosynthates and mineral nutrients, including copper (Udvardi and Poole, 2013; Senovilla et al., 2018).

Legume nodules accumulate high levels of copper (around 50 μg/g) (Senovilla et al., 2018), evidencing an important role for this nutrient in nodule development and/or symbiotic nitrogen fixation. Further evidence is provided by the existence of a nodule-specific copper uptake transporter, MtCOPT1, that is required for optimal nitrogen fixation and cytochrome oxidase activity of bacteroids (Senovilla et al., 2018). Therefore, it should be expected that specific mechanisms are in place to ensure copper allocation to enzymes involved in symbiotic nitrogen fixation in legume nodules, and that this is achieved via dedicated Cu^+^-chaperones. This manuscript provides data supporting this hypothesis, as it reports that *Medtr3g067750* is a nodule-specific Cu^+^-chaperone (NCC) that is essential for symbiotic nitrogen fixation.

## RESULTS

### MtNCC1 is a Cu^+^-chaperone

To identify candidate Cu^+^-chaperones in *Arabidopsis thaliana* and *M. truncatula*, we collected those sequences annotated as such, as well as those annotated as metal-binding domain proteins or HIPP, removing those that did not contain a CXXC Cu^+^-binding motive. Over 30 different candidates were identified in the genomes of these two plant species (Fig. S1). Within *M. truncatula*, there was a nodule-specific candidate, *Medtr3g067750* (NCC1, for Nodule-specific Cu^+^-Chaperone1), as indicated by the available transcriptomic databases (Fig. S2). This expression pattern was validated by qRT-PCR analyses (Fig. 1A). Protein modelling showed the existence of two different protein domains: a N-terminal Atx1-like domain comprised by the first 78 amino acids (MtNCC1_1-78_) that included the CXXC motif, and a C-terminal, intrinsically disordered region, with an E-rich motif (Fig. 1B).

**Figure 1.**
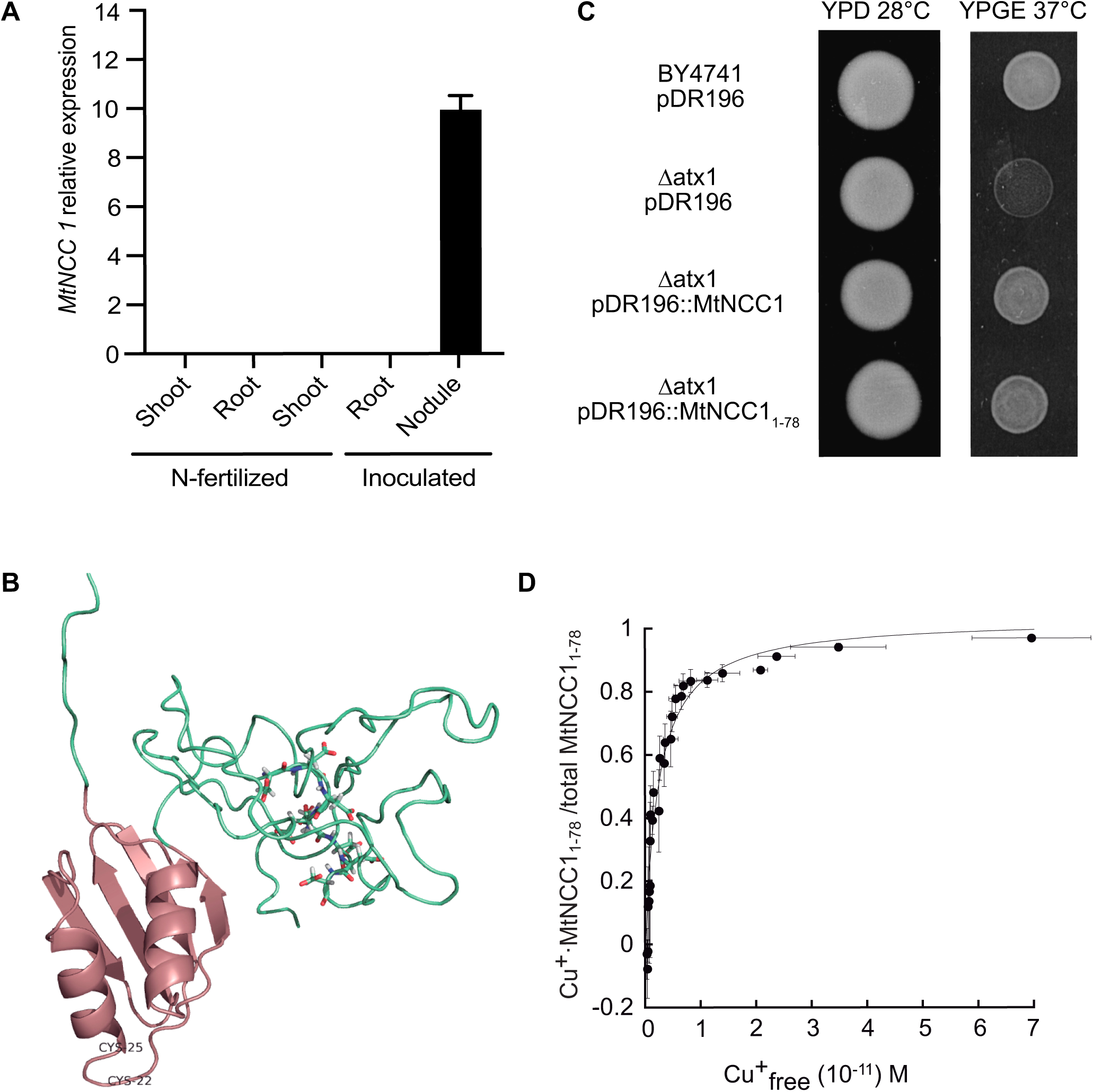
MtNCC1 is a nodule-specific Cu^+^-chaperone. A, *MtNCC1* expression in *S. meliloti-*inoculated (roots, shoots, and nodules) and non-inoculated (roots and shoots) plants relative to standard gene *Ubiquitin carboxyl-terminal hydrolase.* Data are the mean ± SE of three independent experiments with five pooled plants. B, Predicted structure of MtNCC1. The classical Atx1-like domain is indicated in pink, with the two conserved Cu^+^-binding cysteines (C22 and 25). The glutamate-rich region in the C-terminal domain is indicated with wireframes. C, Parental strain BY4741 was transformed with empty pDR196 vector, while the /−atx1 mutant NCC1 was transformed with empty pDR196 or containing MtNCC1 or MtNCC1_1-78_. D, Cu^+^ binding to MtNCC1_1-78_ determined in competition assays with BCA. The data were fit using n = 1.03 ± 0.04 Cu^+^ per protein and K_a_ = 2,45·10^-12^ ± 3.44·10^-13^ M^-1^. Data are the mean ± SE (n = 3).

Yeast complementation assays of the yeast *atx1* mutant were used to determine whether the candidate protein above could function as a Cu^+^-chaperone. This strain cannot grow in non-fermentative carbon sources. However, *atx1* showed wild-type growth when expressing *MtNCC1* or *MtNCC1_1-78_* (Fig. 1C). To further determine whether MtNCC1 could work as a Cu^+^-chaperone, Strep-tagged MtNCC1_1*-78*_ was purified from *Escherichia coli* (Fig. S3) and used to determine Cu^+^-binding stoichiometry and affinity. As expected from an Atx1-like chaperone, MtNCC1_1-78_ bound one Cu^+^ per molecule with an affinity constant of 2.45 pM^-1^ (Fig. 1D). A similar stoichiometry was observed after incubating MtNCC1_1-78_ with a 10-fold molar excess of Cu^+^ and removing the unbound metal (Fig. S4).

### MtNCC1 is located in the cytosol and nucleus of nodule cells

To determine whether MtNCC1 was produced in the nodule, promoter-GUS fusions were transformed in *M. truncatula* plants. As shown in Fig 2A, *MtNCC1* was highly expressed in the infection-differentiation zone (ZII), interzone (IZ) and early fixation zone (ZIII) of the nodules, although lower expression levels could be found in older fixation zone areas. This expression pattern is consistent with the transcriptomic data obtained from laser-captured micro-dissected cells in the Symbimics database (Fig. 2B). Moreover, immunolocalization of hemagglutinin epitope (HA)-tagged MtNCC1 showed a similar distribution (Fig 2C). Higher magnification images of nodule cells showed MtNCC1-HA in the cytoplasm and in the nuclei in infected and uninfected cells (Fig. 2D). This distribution pattern was not observed in un-transformed plants (Fig. S5). Additionally, MtNCC1_1-78_-HA had the same distribution pattern (Fig. S6). Electron-microscopy images using a gold-conjugated antibody revealed that MtNCC1-HA was found in the cytosol, as well as associated with the plasma and symbiosome membranes (Fig. 2E).

**Figure 2.**
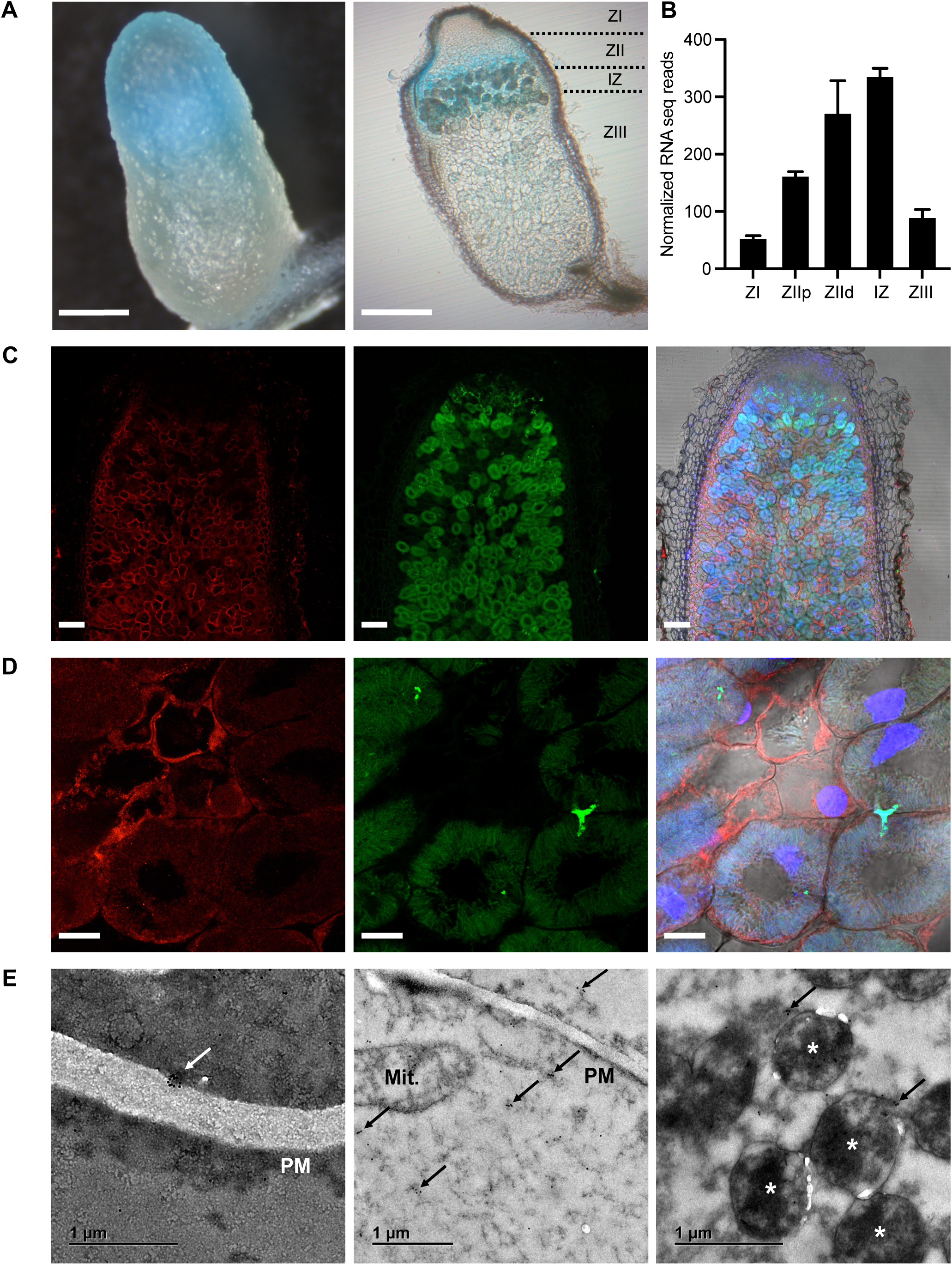
Localization of MtNCC1 in *M. truncatula* nodules. A, GUS staining of 28 dpi nodules expressing the *gus* gene under the control of the *MtNCC1* promoter region. Left panel shows the whole nodule and right panel shows a longitudinal section with the different developmental zones indicated. Bars = 400 μm. B, *MtNCC1* expression in the different nodule zones as indicated in the Symbimics database (http://ia.nt.toulouse.inra.fr/symbimics/). ZI: zone I; ZIIp: zone II proximal; ZIId: zone II distal; IZ: interzone; ZII: zone III. C, Longitudinal section of 28 dpi *M. truncula* nodules expressing *MtNCC1* fused to three HA domains driven by its own promoter. HA-tagged proteins were detected with using an Alexa594 conjugated antibody (red, left panel). Nodules were colonized by a GFP-expressing *S. meliloti* (green, central panel). DNA was stained with DAPI (blue) and overlaid with the previous two channels and the transillumination signal (right panel). Bars = 100 μm. D, Closer view of rhizobia-infected cell in zone II. Bars = 10 μm. E, Immunolocalization of MtNCC1-HA in 28 dpi nodules using gold-conjugated antibodies and electron transmission microscopy. Gold particles are indicated by arrows and asterisks represent bacteroids. PM indicates Plasma membrane, and Mit refers to mitochondria. Bars = 1μm.

### MtNCC1 is required symbiotic nitrogen fixation

To determine the physiological role of MtNCC1, an insertional *ncc1* mutant was obtained from a *M. truncatula Tnt1* mutant collection. This line had an insertion in position +1044 that led to loss of *MtNCC1* expression (Fig. 3A). As expected, the mutant line did not present any apparent phenotype under non-symbiotic conditions (Fig S7), but showed a significantly reduced growth and biomass production in symbiosis (Fig. 3B,C). No significant changes in nodule development were observed compared to wild-type plants (Fig. S8). The nitrogenase activity in *ncc1* plants was severely reduced (Fig. 3D). Analyses of *MtNCC1* wild-type *ncc1* segregants suggested that no additional *Tnt1* insertions elsewhere in the genome could explain these phenotypical effects (Fig. S9). Furthermore, growth and nitrogenase activity were restored by introducing a wild-type copy of *MtNCC1* into the mutant line (Fig 3B-D). Finally, cytochrome oxidase activity was significantly reduced in bacteroids, but to a much lower extent than the nitrogenase activity (Fig. 3E).

**Figure 3.**
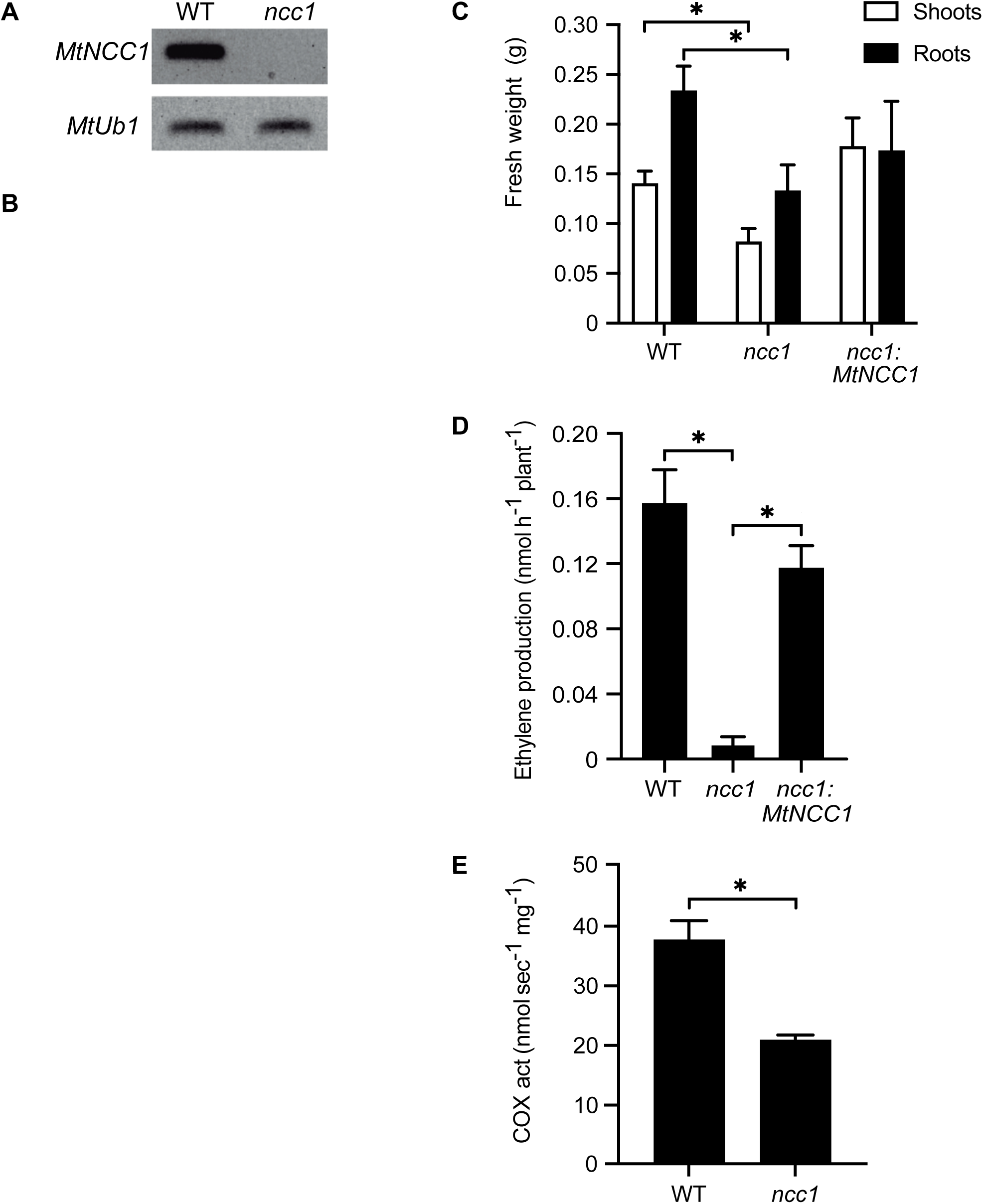
MtNCC1 is required for nitrogen fixation. A, Amplification of *MtNCC1* transcript by RT-PCR in 28 dpi nodules of wild type and *ncc1* mutants. *MtUb1* (*Ubiquitin carboxyl-terminal hydrolase 1*) was used as a constitutive gene. B, Growth of representative plants of wild type (WT), *ncc1* mutant and *ncc1* mutants transformed with *MtNCC1* regulated by its own promoter. Bar = 1cm. C, Fresh weight of shoots and roots of WT, *ncc1* and *ncc1* transformed with *MtNCC1* regulated by its own promoter. Data are the mean ± SE (n = 15). D, Acetylene reduction assay in 28 dpi nodules from WT, *ncc1* mutants and *ncc1* transformed with *MtNCC1* regulated by its own promoter. Data are the mean ± SE (n = 7-13). E, Cytochrome oxidase (COX) activity in bacteroids isolated from 28 dpi nodules of WT and *ncc1* mutant plants. Data are the mean ± SE of at least three sets of 35-40 pooled plants each. Student’s *t*-test was used for statistical analysis (* = P≤ 0.05).

### *MtNCC1* mutation affects a subset of the copper-proteome

Consistent with the role of MtNCC1 in copper metabolism, increasing copper concentrations by 10-fold in the nutrient solution was sufficient to restore wild-type growth and nitrogenase activity in *ncc1* plants (Fig. 4). However, overall copper content in the analysed plant organs (roots, shoots, and nodules) was not significantly altered in *ncc1* plants (Fig 5A). Iron levels were not affected either (Fig. 5B). Importantly, metalloproteomic analyses on nodule soluble protein extracts from wild-type and *ncc1* nodules showed reduced copper contents in a subset of the proteome (Fig. 6A). The largest differences were found around 110 kDa and 60 kDa. As this was a separation only by SEC, the copper proteins only represented a small proportion of the total proteins eluting during the times when a maximal difference between the wild-type and *ncc1* samples was observed. This becomes clear when looking at the profiles of other metals at the same time and comparing the UV/VIS spectra with the metal chromatograms (regions of interest in Figures 6B, 6C, complete time range in Figure S10).

**Figure 4.**
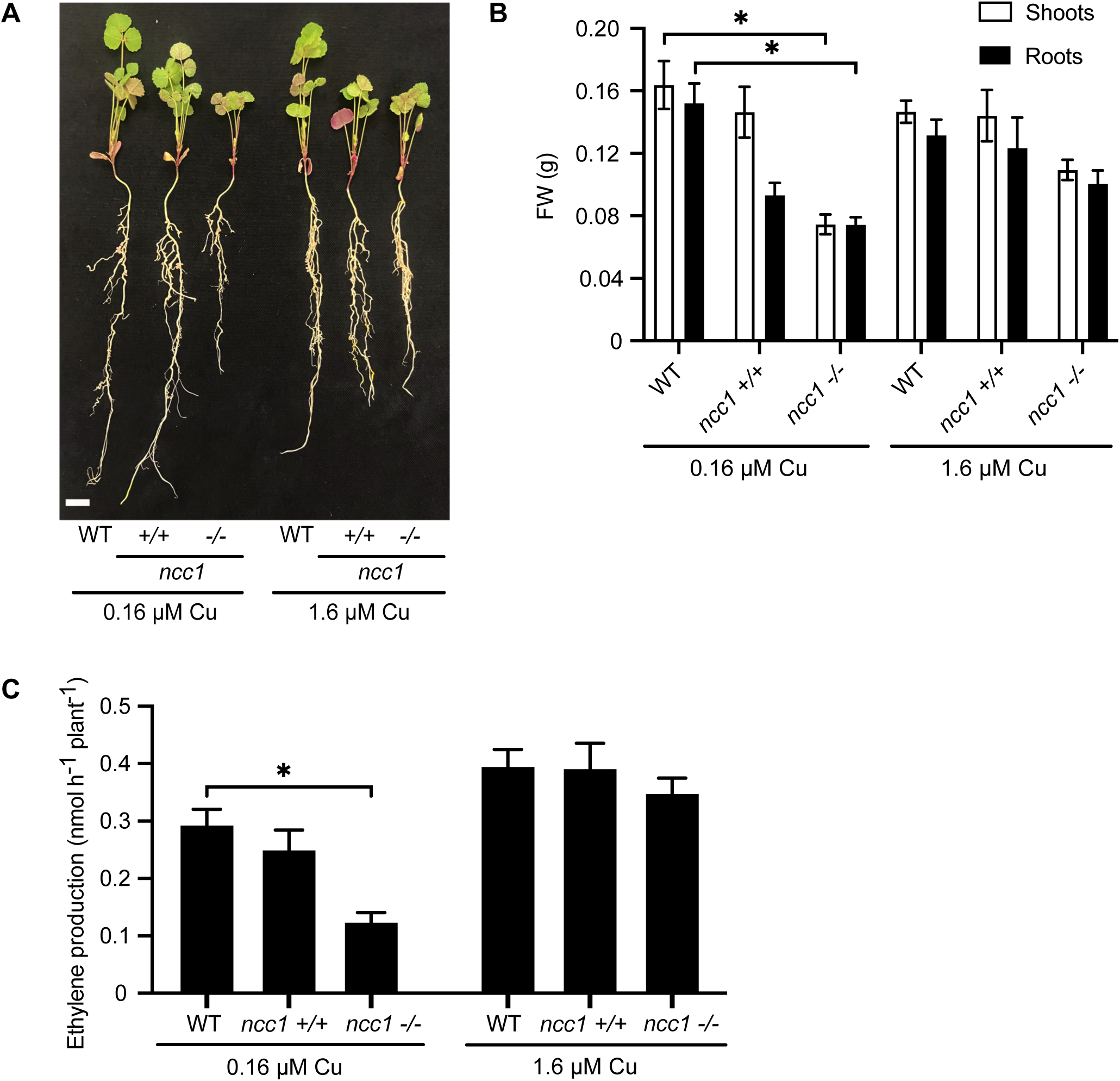
Copper supplementation complements the *ncc1* phenotype. A, Growth of representative plants of wild type (WT) and *ncc1* mutant watered with standard copper concentrations (0.16 μM) and copper excess (1.6 μM). Bar = 1cm. B, Fresh weight of shoots and roots of WT and *ncc1* mutant watered with standard copper concentrations (0.16 μM) and copper excess (1.6 μM). Data are the mean ± SE (n = 16-23). C, Acetylene reduction assay in 28 dpi nodules from WT and *ncc1* mutant watered with standard copper concentrations (0.16 μM) and copper excess (1.6 μM). Data are the mean ± SE (n = 16-23). All comparison were done to WT samples using Student’s *t*-test (* = P≤ 0.05).

**Figure 5.**
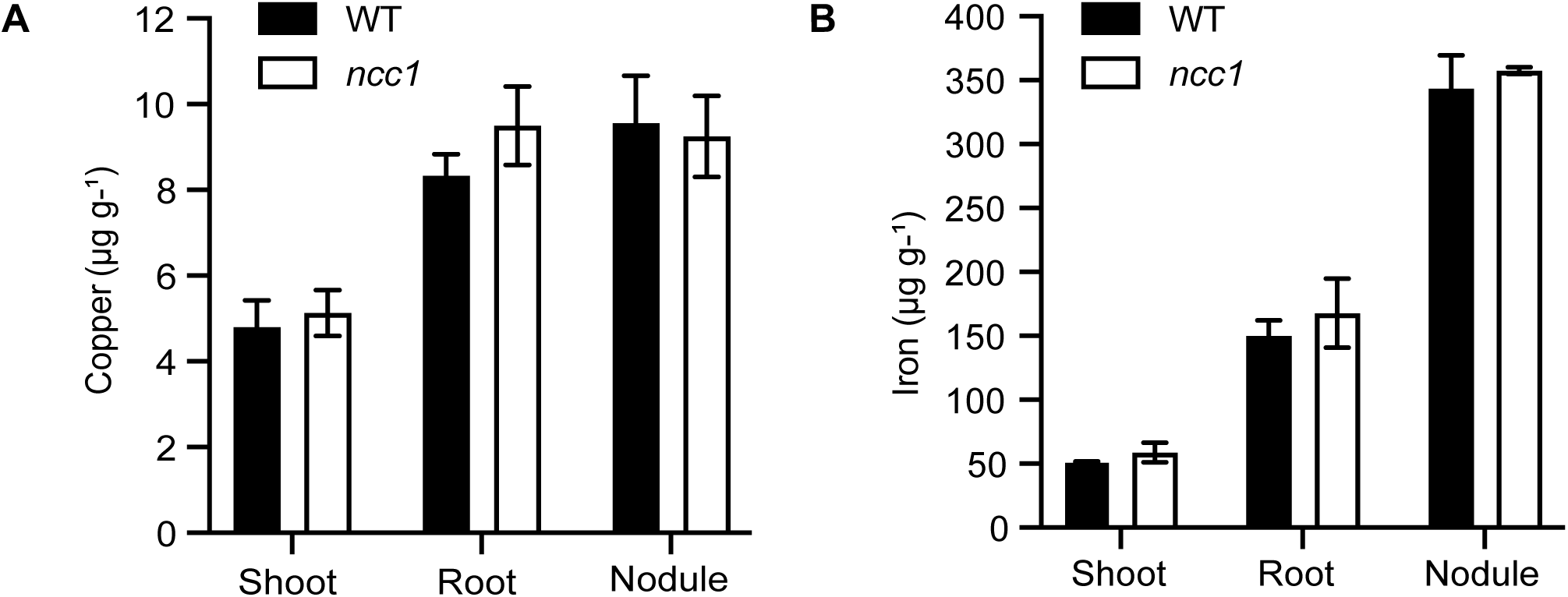
Mutation of *MtNCC1* does not alter the nodule metallome. A, Copper content in 28 dpi shoots, roots and nodules of wild type (WT) and *ncc1* plants. Data are the mean ± SE of three sets of ten pooled plants. B, Iron content in 28 dpi shoots, roots and nodules of wild type (WT) and *ncc1* plants. Data are the mean ± SE of three sets of ten pooled plants.

**Figure 6.**
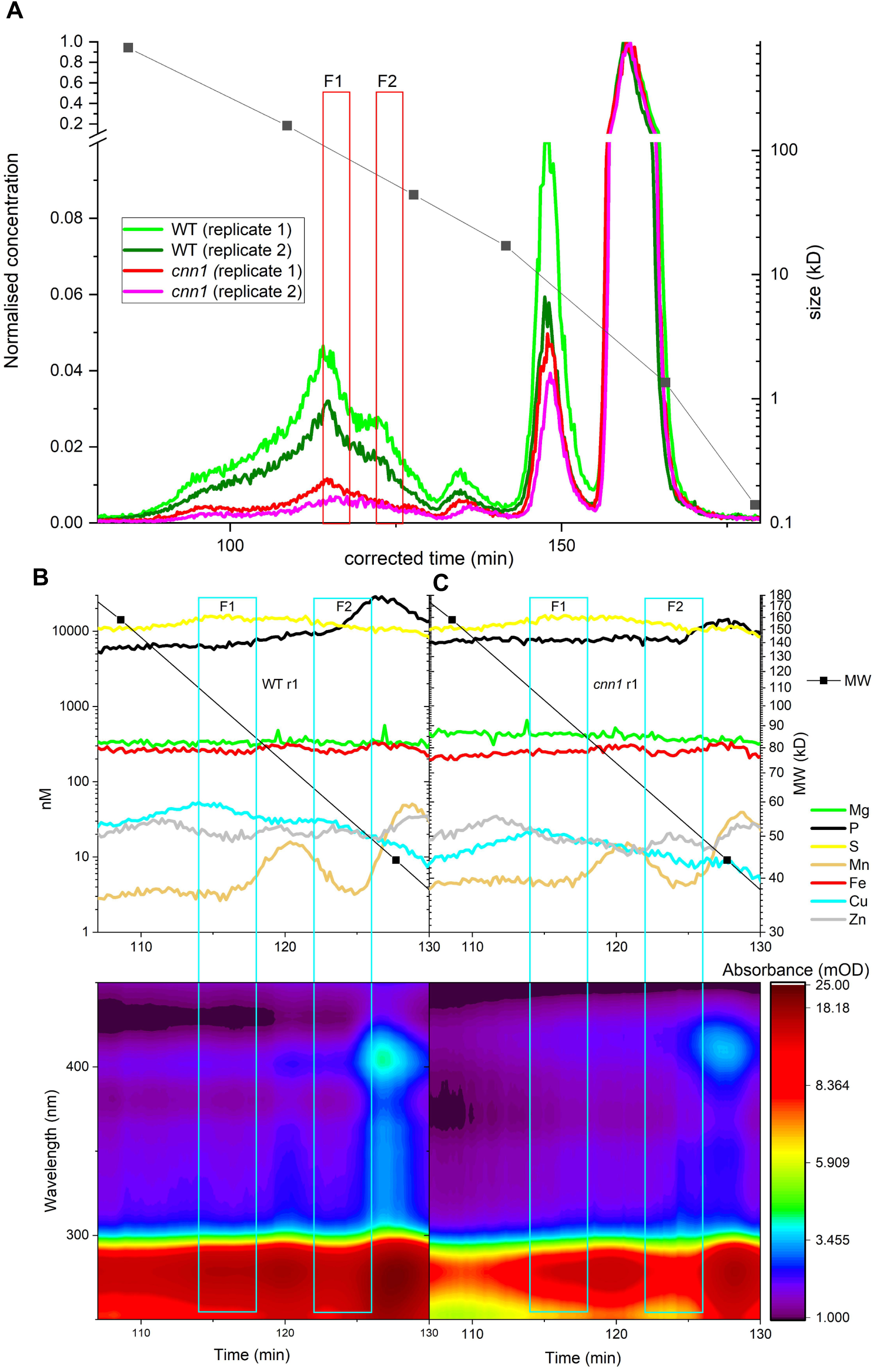
Mutation of *MtNCC1* alters the nodule copper-proteome. A, Metalloproteomics by HPLC-ICPsfMS. The normalized chromatograms show copper content in two independent biological replicates of 28 dpi WT and *ncc1* nodules. Fractions subjected to proteomic analyses are boxed and indicated as F1 and F2. B and C, Comparison of the most important element chromatograms and UV/VIS absorption DAD in the chromatogram region where the biggest differences between the mutant had been seen in the copper chromatogram. B, replicate 1 of the WT, C, replicate 1 of the *ncc1* mutant.

Therefore, although copper-binding proteins were affected, the total number of proteins contained in two fractions (chosen in the MW ranges of max. differences in copper, Fig 6A) in wild type and *ncc1* nodules remained largely the same (Tables S1 and S2); only 70 out of 1529 proteins in fraction 1 and 12 out of 1900 proteins in fraction 2 missing in *ncc1* compared to wild type. None of the ten most abundant of these differentially expressed proteins have a known role in copper-homeostasis (Table 1). Looking at the entire HPLC-ICPsfMS chromatograms in more detail, (Fig S10) reveals additional changes in the metalloproteome that would result from the cross-talk between copper homeostasis and that of other elements. Analysing these changes would be a topic for another study.

**Table 1.** Top ten most abundant proteins detected in wild type replicates and not in ncc1 replicates in Fractions 1 or 2 of the metalloproteomic analyses of nodule soluble proteins.

| Uniprot<br>Accession | Organism | Description | Fraction |
| --- | --- | --- | --- |
| Q4H1G3 | *M. truncatula* | S-adenosylmethionine synthase | 1 |
| A0A072U7P3 | *M. truncatula* | Heat shock cognate 70 kDa protein | 1 |
| P47923 | *M. truncatula* | Nucleoside diphosphate kinase 2 | 1 |
| Q1SL05 | *M. truncatula* | Putative small GTPase superfamily, P-loop<br>containing nucleoside triphosphate hydrolase | 2 |
| G7IEZ9 | *M. truncatula* | Chalcone-flavonone isomerase family protein 2 | 1 |
| Q92M81 | *S. meliloti* | Glyceraldehyde-3-phosphate dehydrogenase | 1 |
| A0A072UDY2 | *M. truncatula* | Presequence protease | 1 |
| Q92SW9 | *S. meliloti* | Recombination protein Rec | 2 |
| Q92RK3 | *S. meliloti* | 4-phospho-D-erythronate dehydrogenase | 1 |
| Q92RH2 | *S. meliloti* | 3,4-dihydroxy-2-butanone 4-phosphate synthase | 1 |

In a further attempt to identify proteins that would accept Cu^+^ by interacting with MtNCC1, Cu^+^-loaded MtNCC1_1-78_ (MtNCC1_1-78_·Cu^+^) was used as a bait for pull-down assays. In order to ensure that the identified proteins were those that specifically interacted with MtNCC1_1-78_·Cu^+^ and not with the resin, control pull-down assays in which no bait was bound to the resin were done in parallel. The proteins specifically retained by MtNCC1_1-78_·Cu^+^ are indicated in Table S3. Approximately, one third of these proteins were also present in the selected fractions from the metalloproteomic analyses (Table S4), and mostly they are putatively involved in D-gluconate catabolism, S-adenosylmethione (SAM) cycle, endoplasmatic reticulum unfolded protein response, and protein import into the nucleus. To further confirm some of these interactions, Bi-molecular Fluorescence Complementation (BiFC) studies were carried out in agroinfiltrated tobacco leaves expressing *MtNCC1_1-78_* together with selected candidates identified in the pull-down experiments. As shown in Fig. 7, MtNCC11_-78_ interacted with thioredoxin-dependent peroxiredoxin (*Medtr7g105830)*, SAM synthase (*Medtr2g046710),* putative universal stress protein (*Medtr1g088640)*, and pathogenesis-related protein (*Medtr2g076010)*. No signal was observed when MtNCC1_1-78_ was expressed in leaves infiltrated with the empty vectors (Fig. S11).

**Figure 7.**
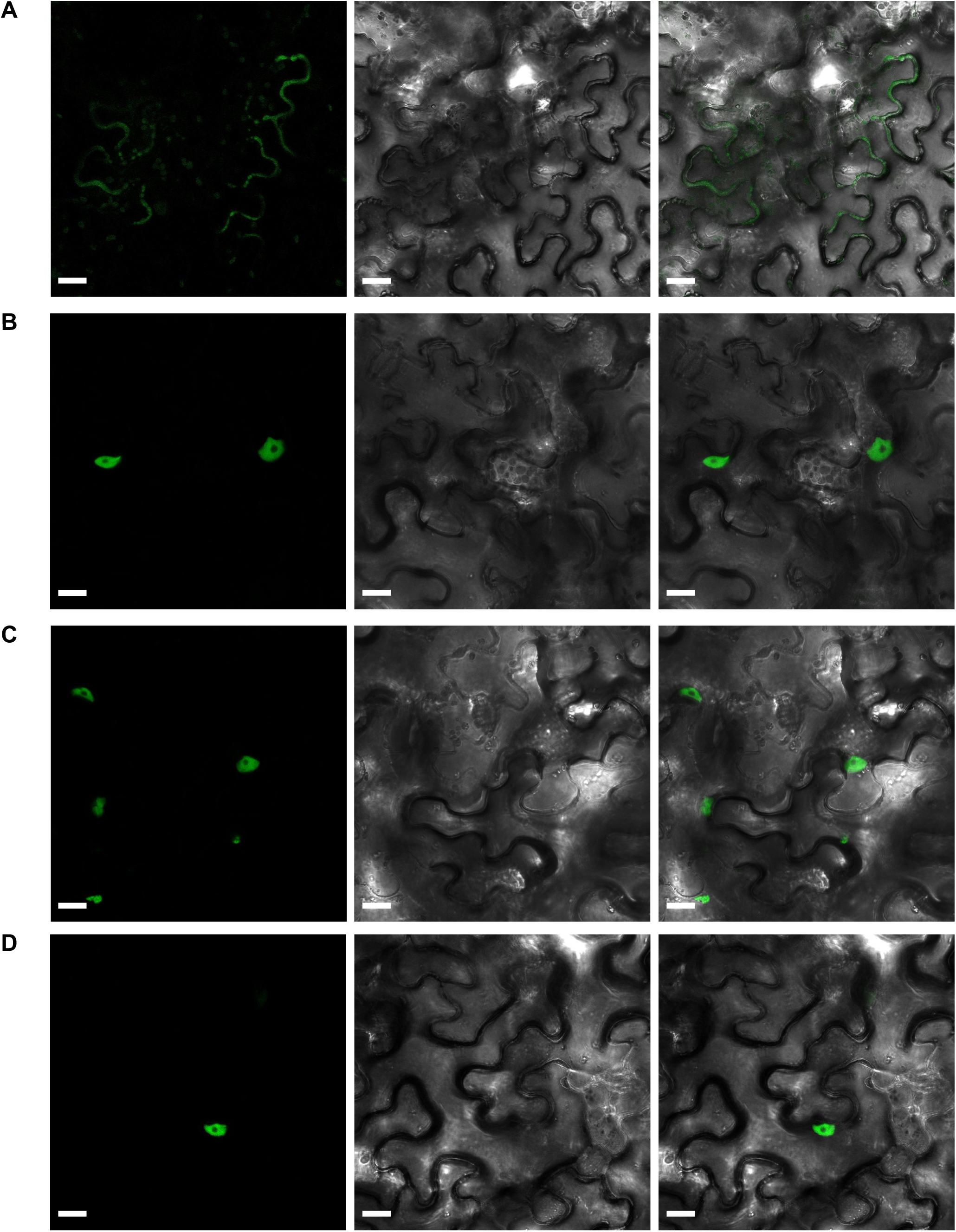
Bimolecular Fluorescent Complementation assays of MtNCC1_1-78_ interactions. Transient co-expression of MtNCC1_1-78_ in pNXGW and candidate proteins thioredoxin-dependent peroxiredoxin (A) and SAM synthase (B) in pCXGW, putative universal stress protein (C) and pathogenesis-related protein (D) in pXCGW, in *N. benthamiana* leaf cells 3 days-post-agroinfiltration. Left panel corresponds to the fluorescent signal of MtNCC1_1-78_ and corresponding interactor (green); central panel, transillumination; right panel, overlaid fluorescent image and transillumination images. Bars = 20 μm.

## DISCUSSION

Symbiotic nitrogen fixation in legume root nodules is a process that requires of a large transfer of transition elements from the host plant to the root nodules (O’Hara, 2001; González-Guerrero et al., 2016). In recent years, many of the transporters mediating this transfer have been identified (Tejada-Jiménez et al., 2015; Abreu et al., 2017; Tejada-Jiménez et al., 2017; Senovilla et al., 2018; Castro-Rodríguez et al., 2020; Escudero et al., 2020), as well as metal-chelating molecules that maintain metal solubility in saps and apoplast (Kryvoruchko et al., 2018; Escudero et al., 2020b). In the case of copper, there is a high degree of specificity in nodules. Host plants, such as *M. truncatula*, express a nodule-specific Cu^+^-uptake transporter, MtCOPT1, to ensure copper uptake by infected cells (Senovilla et al., 2018). Bacteroids also synthesize nodule-specific copper transporters, such as FixI, a P_1B_-ATPase responsible for providing copper for a nodule-specific high-affinity cytochrome oxidase (Kahn et al., 1989; Preisig et al., 1996). However, an additional level of control based on Cu^+^-chaperones must exist so that copper is specifically delivered from very few copper transporters to multiple different copper-enzymes. We have identified MtNCC1 as a representative of this layer of control. The relationship of MtCOPT1 and MtNCC1 roles in nodule copper homeostasis is suggested by the presence of both proteins in the same nodule zones (mainly from late infection to early fixation zones). This hints at a functional pairing between the two proteins, *i.e.* the transporter transfers copper to the chaperone. Moreover, MtNCC1 is not the only copper delivery system to bacteroids. As for *copt1-1*, copper fortification of the nutrient solution is sufficient to restore wild-type growth and nitrogenase activity, suggesting that MtNCC1 is only necessary below a certain copper concentration threshold and that other lower-affinity system(s) can functionally complement MtNCC1.

The connection between the reduction of nitrogenase activity and mutation of *MtNCC1* is not direct. Nitrogenase does not use copper as a cofactor, nor is copper known to be directly needed for its synthesis (Burén et al., 2020). In previous studies (Senovilla et al., 2018), reduction of nitrogenase activity could be explained as the consequence of the reduction of cytochrome oxidase activity that would limit the energy available to bacteroids. However, the partial reduction in cytochrome oxidase activity cannot explain the drastic loss of nitrogenase activity in *ncc1*. The explanation cannot be altered nodule morphology either (wild-type and *ncc1* nodules do not seem different), nor can it be a collateral effect on iron metabolism in the nodule. The reason must instead lie in the subset of proteins that interact with MtNCC1. Many of them, when forming complexes with MtNCC1_1-78_ can be detected in the nuclei. Some of them could be involved in epigenetic regulation (SAM synthases) (Liu et al., 2020), in the response to plant-microbe interactions (pathogenesis-related proteins) (Kaur et al., 2017), or in the coordination between host and bacteroid (Nodule-specific Cysteine-rich peptides, NCRs) (Maróti et al., 2015), but their specific roles in symbiotic nitrogen fixation still remain to be determined. Our metalloproteomic and pull-down assays have unveiled a number of new putative copper-proteins that have to be further validated biochemically and in the context of symbiotic nitrogen fixation. Other candidate copper-proteins might emerge from studying the interaction with the C-terminal domain of MtNCC1. However, all our efforts in that direction were fruitless because of our failure to recover either full MtNCC1 or C-domain MtNCC1 from the inclusion bodies that formed when they were expressed in *E. coli*.

Interestingly, MtNCC1 is not only located in the cytosol or in the proximity of the membranes, it can also be detected in nuclei, as was the case of CCP in Arabidopsis (Chai et al., 2020). However, no nuclear localization signal could be found in the MtNCC1 sequence. Alternatively, the observed nuclear localization could be the result of interactions with other proteins. BiFC studies indicate that this is the case, as only some of the interactions tested led to MtNCC_1-78_ detection in nuclei. The migration of MtNCC1 from cytosol to nucleus will be likely facilitated by importins (Merkle, 2011), some of which have been detected in the pull-down assays. The physiological relevance of this localization is not evident. Theoretically, copper could be delivered to any proteins in the cytosol and then the newly-metallated proteins could migrate to the nucleus. It could be argued that copper transfer to the nucleus could be conditional to specific environmental cues, or that some sort of ternary complex in the nucleus would be needed. These possibilities will have to be specifically tested in the future. However, one role that nuclear MtNCC1 does not seem to play is to deliver Cu^+^ to a putative copper sensor. The existence of a nodule-specific copper sensor can be inferred from the previous work on MtCOPT1. Lack of copper uptake by nitrogen-fixing cells did not result in nodules containing less copper, but the opposite (Senovilla et al., 2018). This was interpreted as a copper-deficiency signal being sent from nodule cells, that led to more copper being delivered. However, since MtCOPT1 was not present, this copper did not reach the cell cytosol and accumulated in the apoplast. This phenotype, however, was not observed in *ncc1* mutants, in which no significant change in copper contents was observed.

In summary, in this work we have shown the existence of a nodule-specific Cu^+^-chaperone that is required for nitrogen fixation. We have identified changes in the nodule copper-proteome as a result of losing MtNCC1 activity, as well a new putative copper-proteins that might play a role in symbiotic nitrogen fixation. Future work will be directed to determining the physiological role of these proteins and to verifying that they can accept copper from MtNCC1.

## METHODS

### Biological materials and growth conditions

*Medicago truncatula* Gaertner R108 seeds were scarified in pure H_2_SO_4_ for 7.5 min. Then, they were washed with cold water and sterilized with 50% bleach for 1.5 min and imbibed in sterile water in darkness overnight. On the following day, seeds were placed on water-agar plates for 48 h at 4°C, and then allowed to germinate at 22°C for 24 h. Seedlings were planted in sterile perlite pots and inoculated with *Sinorhizobium meliloti* 2011 or *S. meliloti* 2011 transformed with pHC60 (Cheng and Walker, 1998), as indicated. Plants were grown in a greenhouse in 16 h light/8 h dark and 22°C and watered with Jenner’s solution or water every 2 d, alternatively (Brito et al., 1994). Nodules were collected at 28 days-post-inoculation (dpi). Non-inoculated plants were cultivated in similar conditions, but they were watered every 2 weeks with Jenner’s solution supplemented with 2 mM NH_4_NO_3_. For hairy-root transformations, *M. truncatula* seedlings were infected with *Agrobacterium rhizogenes* ARqua1 carrying the appropriate binary vector as described (Boisson-Dernier et al., 2001). In agroinfiltration experiments, *Nicotiana benthamiana* (tobacco) leaves were infected with the plasmid constructs in *A. tumefaciens* GV3101 (Deblaere et al., 1985). Tobacco plants were grown under the same conditions as *M. truncatula*.

For yeast complementation assays, *Saccharomyces cerevisiae* strain Δ*atx1* and its parental strain BY4741 (MATa *his3/− 1 leu2/− 0 met15/− 0 ura3/− 0*) were purchased from the Yeast Knockout Collection (GE Dharmacon). Yeasts were grown in yeast peptone dextrose (YPD) or in synthetic dextrose (SD) media supplemented with 2% glucose (Sherman et al., 1983). Phenotypic characterization was performed in yeast peptone ethanol glycerol (YPEG) medium (Li & Kaplan, 2001).

For protein purification, *Escherichia coli* BL21 (DE3) pLysS (Wood, 1966) (*E. coli* str. B F–*ompT gal dcm lon hsdSB (r_B–_m_B–_)* λ(DE3 [*lacI lacUV5-T7p07 ind1 sam7 nin5*]) [*malB^+^*]_K-12_ (λ^S^) pLysS [T7p20 ori_p15A_](Cm^R^) was used to produce the required amount of protein.

### Quantitative real-time RT-PCR

RNA isolation and cDNA synthesis were carried out as previously described (Tejada-Jiménez et al., 2015). Gene expression was studied by quantitative real-time RT-PCR (9700, Applied Biosystems) with primers indicated in Table S5 and normalized to the *M. truncatula Ubiquitin carboxy-terminal hydrolase* gene (*Medtr4g077320*). Real-time cycler conditions have been previously described (González-Guerrero et al., 2010). Determinations were performed with RNA extracted from three independent biological samples, with the threshold cycle determined in triplicate. The relative levels of transcription were determined with the 2^-1′Ct^ method.

### Yeast complementation assays

*MtNCC1* and *MtNCC1_1_*_-78_ cDNAs were cloned into the yeast expression vector pDR196, between PstI and XhoI restriction sites, by homologous recombination in yeast (primers indicated in Table S5). For yeast transformation, a lithium acetate-based method was used as described (Schiestl and Gietz, 1989). Transformants were selected in SD medium by uracil autotrophy. Phenotypic analysis of yeast transformants were done on YPEG plates (Li & Kaplan, 2001).

### Protein Expression and Purification

*MtNCC1_1_*_-*78*_ was obtained by PCR using *MtNCC1* cDNA as the template (primers indicated in Table S5). The resulting cDNA was cloned between *NdeI* and *BamHI* restriction sites in a modified version of pET16b (courtesy of Dr. Luis Rubio). This plasmid adds two N-terminal streptavidin (N-Twin-Strep (N-TS)) tag sequences. Protein expression was induced for 3 h at 37°C by the addition of 1 mM IPTG to *E. coli* BL21 cells (Wood, 1966). Cells expressing soluble MtNCC1_1-78_ were disrupted in 100 mM Tris (pH 8.0), 150 mM NaCl (buffer W) using a French press. Homogenates were centrifuged at 54,000 g for 1 h. The protein was purified with a Strep-Tactin XT 4Flow high-capacity column (IBA lifesciences) and stored in 10% glycerol in buffer W at -80°C. Protein quantification was performed in accordance with Bradford assay (Bradford, 1976).

### Cu^+^-binding affinity

MtNCC1_1-78_·Cu^+^ *K_a_* value was obtained by using a competition assay with BCA followed by colorimetric determination of the BCA_2_·Cu^+^ complex at 360 nm. MtNCC1_1-78_·Cu^+^ *K_a_* was determined by titrating with Cu^+^ in a solution of 20 μM BCA, 100 μM MtNCC1_1-78_ in 50 mM HEPES (pH 7.5), 200 mM NaCl, and 200 μM ascorbate (buffer H). The BCA_2_·Cu^+^ molar extinction coefficient, ε_360_ = 20,600 cm^-1^ M^-1^, was determined by titrating μM Cu^+^ with 0–10 μM BCA in buffer H. Free metal concentrations were calculated from *K*_BCA_ = [BCA_2_·Cu^+^]/[BCA_free_]^2^[Cu^+^_free_], where *K*_BCA_ is the association constant for BCA_2_·Cu^+^ (4.60 x 10^14^ M^-2^) (Yatsunyk and Rosenzweig, 2007; González-Guerrero and Argüello, 2008). The MtNCC_1-78_·Cu^+^ *K_a_* value was calculated by using ϖ = [Cu^+^_free_]*^n^K_a_*/(1 + *K_a_* [Cu^+^_free_]*^n^*), where ϖ is the molar ratio of metal bound to protein and *n* is the number of metal-binding sites. Reported errors for *Ka* and *n* are asymptotic standard errors provided by the fitting software (Origin; OriginLab).

### GUS staining

*MtNCC1* promoter (*MtNCC1p*; 2 kb upstream of the *MtNCC1* start codon) was amplified using the primers indicated in Table S5, then cloned into pDONR207 (Invitrogen) and transferred to pGWB3 vector containing the *GUS* gene (Nakagawa et al., 2007), using Gateway Cloning technology (Invitrogen). *M. truncatula* hairy-roots transformation was carried out as described above. Transformed seedlings were planted in sterile perlite pots and inoculated with *S. meliloti* 2011. GUS activity was measured in nodules of 28 dpi plants as described (Vernoud et al., 2007).

### Immunolocalization

A DNA fragment containing the full-length *MtNCC1* genomic region and the *MtNCC1p* was cloned into the pGWB13 (Nakagawa et al., 2007) by Gateway technology (Invitrogen). This vector fuses three HA epitopes in the C-terminus of the protein. *M. truncatula* hairy-root transformation was carried out as described above (Boisson-Dernier et al., 2001). Transformed seedlings were planted in sterile perlite pots and inoculated with *S. meliloti* 2011 integrating the pHC60 vector that constitutively expresses GFP. 28 dpi nodules were collected and fixed in 4% (w/v) paraformaldehyde, 2.5% (w/v) sucrose in phosphate-buffered saline (PBS) at 4°C overnight. Next day, nodules were washed in PBS, and 100 μm sections were generated with a Vibratome 1000 plus (Vibratome). Then, the sections were dehydrated using a methanol series (30%, 50%, 70% and 100% [v/v] in PBS) for 5 min and then rehydrated. Cell walls were permeabilized with 4% (w/v) cellulase in PBS for 1 h at room temperature and treated with 0.1% (v/v) Tween 20 in PBS for 15 min. The sections were blocked with 5% (w/v) bovine serum albumin (BSA) in PBS and incubated then with an anti-HA mouse monoclonal antibody (Sigma) for 2 h at room temperature. After washing the primary antibody, the sections were incubated with an Alexa594-conjugated anti-mouse rabbit monoclonal antibody (Sigma) for 1 h at room temperature. After removing the unbound secondary antibody, DAPI (40,6-diamidino-2-phenylindole) was used to stain the DNA. Images were acquired with a confocal laser-scanning microscope (Leica SP8) using excitation lights at 488 nm for GFP and at 561 nm for Alexa 594.

For gold-immunolocalization, 28 dpi nodules were collected and fixed in 1% formaldehyde and 0.5% glutaraldehyde in 50 mM potassium phosphate (pH 7.4) for 2 h. The fixation solution was renewed and incubated for an additional 1.5 h. Samples were washed in 50 mM potassium phosphate (pH 7.4) 3 times during 30 min and 3 times for 10 min. Nodules were dehydrated by incubating with ethanol dilution series of 30%, 50%, 70%, and 90% during 10 min, 96% for 30 min, and 100% for 1 h. Samples were incubated with a series of ethanol and LR-white resin (London Resin Company Ltd, UK) dilutions: 1:3 for 3 h, 1:1 overnight, and 3:1 for 3 h, and left in LR-white resin for 48 h. All incubations were done at 4°C. Nodules were placed in gelatine capsules previously filled with LR-white resin and polymerized at 60 °C for 24 h. Ultra-thin sections were cut at Centro Nacional de Microscopia Electrónica (Universidad Complutense de Madrid, Spain) with Reichert Ultracut S-ultramicrotome fitted with a diamond knife. The sections were blocked in 2% bovine serum albumin (BSA) in PBS for 30 min. An anti-HA rabbit monoclonal antibody (Sigma) was used (dilution 1:20 in PBS) as primary antibody. The samples were washed 10 times in PBS for 2 min. As a secondary antibody, 1:150 dilution in PBS of antirabbit goat antibody conjugated to a 15 nm gold particle (BBI solutions) was used. Incubation was performed for 1 h. The sections were washed 10 times with PBS for 2 min and 15 times with water for 2 min, stained with 2% uranyl acetate, and visualized in a JEM 1400 electron microscope at 80 kV.

### Acetylene reduction assay

Nitrogenase activity was determined by the acetylene reduction assay (Hardy et al., 1968). Nitrogen fixation was analyzed in mutant and control plants at 28 dpi in 30-ml vials fitted with rubber stoppers. Each tube contained one independently transformed plant. Three ml of air inside the vial was replaced by 3 ml of acetylene and then they were incubated at room temperature for 30 min. Gas samples (0.5 ml) were analyzed in a Shimadzu GC-8A gas chromatograph fitted with a Porapak N column. The amount of ethylene produced was determined by measuring the height of the ethylene peak relative to the background. After measurements, nodules were collected from roots for weighing.

### Cytochrome oxidase activity

Nodules from 28 dpi plants were collected and used for bacteroid isolation, as described by (Brito et al., 1994) with modifications. Nodules from 50 plants (0.1-0.3 g nodules) were pooled together and crushed with 33% (w/w) of polyvinylpyrrolidone and 1 ml of extraction buffer (38 mM K_2_HPO_4_, 24 mM KH_2_PO_4_, 2.4 mM MgCl_2_). Three consecutive centrifugations were performed with the supernatant of the previous one, two at 1000 g (1 and 5 min) and a final one at 5000 g for 10 min. The final pellet was resuspended in 500 μL of resuspension buffer (50 mM HEPES, 200 mM NaCl pH 7). Cytochrome oxidase activity was determined by N,N,N’,N’-tetramethyl-p-phenylenediamine (TMPD) oxidation assay. The reaction was started by adding TMPD to the bacteroid sample at a final concentration of 2.7 mM. To obtain the reaction kinetics, each sample was measured at OD_520_ each 10 s for 3 min. To measure protein content and to calculate specific activity, the bacteroid suspension was lysed in 10% SDS at 90°C for 5 min. The amount of protein was determined with the Pierce^TM^ BCA Protein Assay Kit (Thermo Scientific, Waltham, MA, USA).

### Metal content determination

To determine metal content, Inductively Coupled Plasma-Mass Spectrometry (ICP-MS) was performed for three independent sets of 28 dpi roots, shoots and nodules pooled from ten plants. Elemental analysis with ICP-MS was carried out at the Unit of Metal Analysis from the Scientific and Technology Centre, Universidad de Barcelona (Spain). These samples were treated with HNO_3_, H_2_O_2_ and HF in a Teflon reactor at 90°C. The resulting homogenates were diluted with deionized water. Final volumes were calculated by weight and weight: volume ratios. In parallel, samples were digested with three blanks. Metal determination was carried out in an Agilent 7500cw with standard instrument conditions. Calibration and internal standardization for obtaining the calibration curve was done with five solutions prepared from certified NIST standards

### Metalloproteomics

Metalloproteomic analyses were performed on 150 28 dpi nodules from wild type or *ncc1* plants. Soluble and membrane proteins were extracted from the nodules as described by Andresen et al. (2016). Afterwards, proteins were separated by size exclusion chromatography coupled to ICP-sfMS as described (Küpper et al., 2019). Briefly, two Superdex Increase 200 10 x 300 mm and one Superdex Increase 75 10 x 300 mm size exclusion columns were used to separate the nodule protein extracts in 150 mM ammonium bicarbonate buffer with 0.2 mM DDM. Detection was achieved with a customized sector field ICP-MS (Element XR-2 with jet interface and desolvating injection, Thermo Scientific, Bremen Germany) and a diode array detector coupled to a metal-free HPLC system (Azury system. Knauer, Germny). Metal-EDTA complexes in the same buffer as the protein extracts were used to calibrate concentration determination. A gel filtration calibration standard (Bio-rad with added PABA) was used to determine size and molecular weight. Calibration of metal concentrations in the ICP-sfMS chromatograms was performed as described in Küpper et al. (2019). Proteins were identified in the Proteomic Unit of Universidad Complutense de Madrid (Spain).

### Proteomics

Proteomic analyses were performed in the Proteomics Unit of Complutense University of Madrid, a member of Proteored. Samples were liophylized and resuspended in 100 μl of 25 mM ammonium bicarbonate buffer and quantified. Proteins were reduced in 10 mM DTT at 56°C for 60 min, and subsequently alkylated in 25 mM iodocetamide for 60 min in darkness at room temperature. Samples were digested with Trypsine/LysC protease mix (Pierce, ThermoFisher) at a 1:30 (w/w) ration overnight at 37°C. Desalted and concentrated peptides were liophylized and reconstituted in 15 μl 2% acetonitrile, 0.1% formic acid. One μg of peptides from each sample were analysed by liquid nano-chromatography (Vanquish Neo, Thermo Scientific) coupled to a high-resolution mass spectrometer (Q-Exactive HF (Thermo Scientific). The acquired MS/MS spectra were analysed using Proteome Discoverer 3 (Thermo Scientific) using Mascot 2.8 search engine and the databases SwissProt, *M. truncatula*, *S. meliloti* downloaded from Uniprot. “Correctly” identified proteins are those that have a False Discovery Rate (FDR) below 1 % and at least on single peptide identified with high confidence (above 99 %).

### Pull-down assay

28 grams of fresh *M. truncatula* R108 nodules (from approximately 2.500 plants) were homogenized manually with a mortar and pestle. The homogenate was centrifuged at 20,000 g 4°C for 30 min. To separate soluble from membrane proteins, the supernatant (“cell free” extract) was centrifuged at 100,000 g 4°C for 1 h. The resulting supernatant containing soluble proteins was loaded in Strep-Tactin XT 4Flow High-Capacity column (IBA lifesciences) previously saturated with N-TS-MtNCC_1-78_·Cu^+^. Co-eluting proteins were identified by liquid chromatography Mass Spectrometry (LC-MS/MS) at the Proteomic Unit of the Universidad Complutense de Madrid (Spain).

### Bimolecular fluorescence complementation

*MtNCC1_1-78_* CDS was fused at the N-terminus to the N-fragment of YFP (Yellow Fluorescence Protein) in the Gateway vector pNXGW (Kim et al., 2009). The CDS of MtNCC1candidate interactors were fused to the C-fragment of CFP (Cyan Fluorescence Protein) at both N and C-terminus in the Gateway vectors pCXGW and pXCGW (Kim et al., 2009), respectively. Primers for cloning are indicated in Table S5. These constructs were introduced into *A. tumefaciens* GV3101 (Deblaere et al., 1985). *N*. *benthamiana* leaves were infiltrated as previously described (Senovilla et al., 2018). Leaves were examined after 3 d by confocal laser-scanning microscopy (Zeiss LSM 880) with excitation light of 488 nm for GFP.

### Statistical tests

Data were analyzed using Student’s unpaired t-test to calculate statistical significance of observed differences. Test results with P-values lower than 0.05 were considered as statistically significant.

### Data availability

All the data included in this manuscript is available to interested researchers upon request. Proteomics raw data have been deposited in Proteomics Identification Database (PRIDE).

## AUTHOR CONTRIBUTIONS

CN-G performed most of the experimental work in this manuscript. JL-M contributed to the phenotype and confocal immunolocalization work and carried out the GUS, sequence comparison of *A. thaliana* and *M. truncatula* candidate Cu^+^-chaperones, and the yeast complementation studies. HK and SNHB performed and analysed the metalloproteomic studies. MR-S contributed to plasmid generation and plant management. AP-L collaborated in the BiFC studies. JW and KSM produced and characterized the *M. truncatula Tnt1* collection. SB optimized MtNCC1_1-78_ purification. JI, VE, and MG-G analysed data. VE and MG-G were responsible for overall research supervision and prepared the manuscript with input from all other authors.

## ACKNOWLEDGEMENTS

The authors would like to thank Dr. Luis Fernández-Pacios (CBGP, UPM-INIA/CSIC, Spain) for his help with MtNCC1 modelling, and Dr. Luis Oñate (CBGP, UPM-INIA/CSIC, Spain) for giving us the plasmids used for the BiFC assays. We would also like to thank other members of laboratory 279 at Centro de Biotecnología y Genómica de Plantas (UPM-INIA/CSIC) for their support and feedback in preparing this manuscript.

## REFERENCES

1. Abreu, I., Saez, A., Castro-Rodríguez, R., Escudero, V., Rodríguez-Haas, B., Senovilla, M., Larue, C., Grolimund, D., Tejada-Jiménez, M., Imperial, J., et al. (2017). *Medicago truncatula* Zinc-Iron Permease6 provides zinc to rhizobia-infected nodule cells. Plant Cell Environ. 40:2706–2719.

2. Andresen, E., Peiter, E., Küpper, H. (2018). Trace metal metabolism in plants. J Exp Bot 69:909–954.

3. Andresen, E., Kappel, S., Stärk, H.J., Riegger, U., Borovec, J., Mattusch, J., Heinz, A., Schmelzer, C.E.H., Matoušková, Š., Dickinson, B., Küpper, H. (2016). Cadmium toxicity investigated at the physiological and biophysical levels under environmentally relevant conditions using the aquatic model plant *Ceratophyllum demersum* L. New Phytol 210:1244–1258

4. Attallah, C. v., Welchen, E., Martin, A. P., Spinelli, S. v., Bonnard, G., Palatnik, J. F., and Gonzalez, D. H. (2011). Plants contain two SCO proteins that are differentially involved in cytochrome c oxidase function and copper and redox homeostasis. J Exp Bot 62:4281–4294.

5. Blaby-Haas, C. E., Padilla-Benavides, T., Stübe, R., Argüello, J. M., and Merchant, S. S. (2014). Evolution of a plant-specific copper chaperone family for chloroplast copper homeostasis. Proc Nat Acad Sci USA 111:E5480–E5487.

6. Boisson-Dernier, A., Chabaud, M., Garcia, F., Bécard, G., Rosenberg, C., and Barker, D. G. (2001). *Agrobacterium rhizogenes*-transformed roots of *Medicago truncatula* for the study of nitrogen-fixing and endomycorrhizal symbiotic associations. Mol Plant Microbe Interact 14:695–700.

7. Bradford, M. M. (1976). A rapid and sensitive method for the quantitation of microgram quantities of protein utilizing the principle of protein-dye binding. Anal Biochem 7:248–254.

8. Brito, B., Palacios, J. M., Hidalgo, E., Imperial, J., and Ruiz-Argueso, T. (1994a). Nickel availability to pea (*Pisum sativum* L.) plants limits hydrogenase activity of *Rhizobium leguminosarum* bv. *viciae* bacteroids by affecting the processing of the hydrogenase structural subunits. J Bacteriol 176:5297–5303.

9. Burén, S., Jiménez-Vicente, E., Echavarri-Erasun, C., and Rubio, L. M. (2020). Biosynthesis of nitrogenase cofactors. Chem Rev 120:4921–4968.

10. Burkhead, J. L., Gogolin Reynolds, K. A., Abdel-Ghany, S. E., Cohu, C. M., and Pilon, M. (2009). Copper homeostasis. New Phytol. 182:799–816.

11. Castro-Rodríguez, R., Abreu, I., Reguera, M., Novoa-Aponte, L., Mijovilovich, A., Escudero, V., Jiménez-Pastor, F. J., Abadía, J., Wen, J., Mysore, K. S., et al. (2020). *The Medicago truncatula* Yellow Stripe1-Like3 gene is involved in vascular delivery of transition metals to root nodules. J Exp Bot 71:7257–7269.

12. Chai, L.-X., Dong, K., Liu, S.-Y., Zhang, Z., Zhang, X.-P., Tong, X., Zhu, F.-F., Zou, J.-Z., and Wang, X.-B. (2020). A putative nuclear copper chaperone promotes plant immunity in Arabidopsis. J Exp Bot 71:6684–6696.

13. Changella, A. (2003). Molecular basis of metal-ion selectivity and zeptomolar sensitivity by CueR. Science 301:1383–1397.

14. Cheng, H. P., and Walker, G. C. (1998). Succinoglycan is required for initiation and elongation of infection threads during nodulation of alfalfa by *Rhizobium meliloti*. J Bacteriol 180:5183–5191.

15. Chu, C.-C., Lee, W.-C., Guo, W.-Y., Pan, S.-M., Chen, L.-J., Li, H., and Jinn, T.-L. (2005). A copper chaperone for superoxide dismutase that confers three types of copper/zinc Superoxide dismutase activity in Arabidopsis. Plant Physiol 139:425– 436.

16. Deblaere, R., Bytebier, B., de Greve, H., Deboeck, F., Schell, J., van Montagu, M., and Leemans, J. (1985). Efficient octopine Ti plasmid-derived vectors for *Agrobacterium*-mediated gene transfer to plants. Nucleic Acids Res 13:4777.

17. Downie, J. A. (2014). Legume nodulation. Curr. Biol. 24:R184–R190.

18. Escudero, V., Abreu, I., Tejada-Jiménez, M., Rosa-Núñez, E., Quintana, J., Prieto, R. I., Larue, C., Wen, J., Villanova, J., Mysore, K. S., et al. (2020a). *Medicago truncatula* Ferroportin2 mediates iron import into nodule symbiosomes. New Phytologist 228:194–209.

19. Escudero, V., Abreu, I., del Sastre, E., Tejada-Jiménez, M., Larue, C., Novoa-Aponte, L., Castillo-González, J., Wen, J., Mysore, K. S., Abadía, J., et al. (2020b). Nicotianamine Synthase 2 Is required for symbiotic nitrogen fixation in *Medicago truncatula* nodules. Front. Plant Sci 10.

20. Flis, P., Ouerdane, L., Grillet, L., Curie, C., Mari, S., and Lobinski, R. (2016). Inventory of metal complexes circulating in plant fluids: a reliable method based on HPLC coupled with dual elemental and high-resolution molecular mass spectrometric detection. New Phytol. 211:1129–1141.

21. Goldstein, S., Meyerstein, D., and Czapski, G. (1993). The Fenton reagents. Free Radic Biol Med 15:435–445.

22. González-Guerrero, M., and Argüello, J. M. (2008). Mechanism of Cu^+^-transporting ATPases: Soluble Cu^+^ chaperones directly transfer Cu^+^ to transmembrane transport sites. Proc Natl Acad Sci U S A 105:5992–5997.

23. González-Guerrero, M., Raimunda, D., Cheng, X., and Argüello, J. M. (2010). Distinct functional roles of homologous Cu^+^ efflux ATPases in Pseudomonas aeruginosa. Mol Microbiol 78:1246–1258.

24. González-Guerrero, M., Escudero, V., Sáez, Á., and Tejada-Jiménez, M. (2016). Transition metal transport in plants and associated endosymbionts. Arbuscular mycorrhizal fungi and rhizobia. Front. Plant Sci. 7:1088.

25. Hardy, R. W. F., Holsten, R. D., Jackson, E. K., and Burns, R. C. (1968). The acetylene-ethylene assay for N_2_ Fixation: Laboratory and field evaluation. Plant Physiol 43:1185–1207.

26. Johnston, A. W., Yeoman, K. H., and Wexler, M. (2001). Metals and the rhizobial-legume symbiosis - uptake, utilization and signalling. Adv. Microb. Physiol 45:113– 156.

27. Kahn, D., David, M., Domergue, O., Daveran, M. L., Ghai, J., Hirsch, P. R., and Batut, J. (1989). *Rhizobium meliloti fixGHI* sequence predicts involvement of a specific cation pump in symbiotic nitrogen fixation. J Bacteriol 171:929–939.

28. Kaur, A., Pati, P. K., Pati, A. M., and Nagpal, A. K. (2017). In-silico analysis of cis-acting regulatory elements of pathogenesis-related proteins of *Arabidopsis thaliana* and *Oryza sativa*. PLoS One 12:e0184523.

29. Kim, J. G., Li, X., Roden, J. A., Taylor, K. W., Aakre, C. D., Su, B., Lalonde, S., Kirik, A., Chen, Y., Baranage, G., et al. (2009). *Xanthomonas* T3S effector XopN suppresses PAMP-triggered immunity and interacts with a tomato atypical Receptor-Like Kinase and TFT1. Plant Cell 21:1305–1323.

30. Kryvoruchko, I. S., Routray, P., Sinharoy, S., Torres-Jerez, I., Tejada-Jiménez, M., Finney, L. A., Nakashima, J., Pislariu, C. I., Benedito, V. A., González-Guerrero, M., et al. (2018). An iron-activated citrate transporter, MtMATE67, is required for symbiotic nitrogen fixation. Plant Physiol 176:2315–2329.

31. Küpper, H., Küpper, F., Spiller, M. (1996). Environmental relevance of heavy metal substituted chlorophylls using the example of submersed water plants. J Exp Bot 47:259–266.

32. Küpper, H., Šetlík, I., Spiller, M., Küpper F.C., Prášil, O. (2002). Heavy metal-induced inhibition of photosynthesis: targets of in vivo heavy metal chlorophyll formation. J Phycol 38:429–441.

33. Küpper, H., Andresen, E. (2016). Mechanisms of metal toxicity in plants. Metallomics 8:269–285.

34. Küpper, H., Bokhari, S. N. H., Jaime-Pérez, N., Lyubenova, L., Ashraf, N., and Andresen, E. (2019). Ultratrace metal speciation analysis by coupling of sector-field ICP-MS to high-resolution size exclusion and reversed-phase liquid chromatography. Anal Chem 91:10961–10969.

35. Li, L., and Kaplan, J. (2001). The yeast gene MSC2, a member of the Cation Diffusion Facilitator family, affects the cellular distribution of zinc. J Biol Chem 276:5036– 5043.

36. Lin, S.-J., Pufahl, R. A., Dancis, A., O’Halloran, T. v, and Culotta, V. C. (1997). A role for the *Saccharomyces cerevisiae* ATX1 gene in copper trafficking and iron transport. J Biol Chem 272:9215–9220.

37. Liu, M., Saha, N., Gajan, A., Saadat, N., Gupta, S. v., and Pile, L. A. (2020). A complex interplay between SAM synthetase and the epigenetic regulator SIN3 controls metabolism and transcription. J Biol Chem 295:375–389.

38. Macomber, L., and Imlay, J. A. (2009). The iron-sulfur clusters of dehydratases are primary intracellular targets of copper toxicity. Proc. Natl. Acad. Sci. U S A 106:8344–8349.

39. Maróti, G., Downie, J. A., and Kondorosi, E. (2015). Plant cysteine-rich peptides that inhibit pathogen growth and control rhizobial differentiation in legume nodules. Curr Opin Plant Biol 26:57–63.

40. Marschner, H., and Marschner, P. (2011). Marschner’s Mineral Nutrition of Higher Plants. Elsevier Science.

41. Merkle, T. (2011). Nucleo-cytoplasmic transport of proteins and RNA in plants. Plant Cell Rep 30:153–176.

42. Mira, H., Martínez-García, F., and Peñarrubia, L. (2001). Evidence for the plant-specific intercellular transport of the Arabidopsis copper chaperone CCH. Plant J. 25:521–528.

43. Nakagawa, T., Kurose, T., Hino, T., Tanaka, K., Kawamukai, M., Niwa, Y., Toyooka, K., Matsuoka, K., Jinbo, T., and Kimura, T. (2007). Development of series of gateway binary vectors, pGWBs, for realizing efficient construction of fusion genes for plant transformation. J Biosci Bioeng 104:34–41.

44. O’Hara, G. W. (2001). Nutritional constraints on root nodule bacteria affecting symbiotic nitrogen fixation: a review. Aust. J. Exp. Agr. 41:417–433.

45. Palumaa, P., Kangur, L., Voronova, A., and Sillard, R. (2004). Metal-binding mechanism of Cox17, a copper chaperone for cytochrome c oxidase. Biochem J 382:307–314.

46. Preisig, O., Zufferey, R., Thony-Meyer, L., Appleby, C. A., and Hennecke, H. (1996). A high-affinity cbb3-type cytochrome oxidase terminates the symbiosis-specific respiratory chain of *Bradyrhizobium japonicum*. J. Bacteriol. 178:1532–1538.

47. Rae, T. D., Schmidt, P. J., Pufahl, R. A., Culotta, V. C., and v. O’Halloran, T. (1999). Undetectable intracellular free copper: The requirement of a copper chaperone for superoxide dismutase. Science 284:805–808.

48. Robinson, N. J., and Winge, D. R. (2010). Copper metallochaperones. Annu. Rev. Biochem. 79:537–562.

49. Schiestl, R. H., and Gietz, R. D. (1989). High efficiency transformation of intact yeast cells using single stranded nucleic acids as a carrier. Curr Genet 16:339–346.

50. Senovilla, M., Castro-Rodríguez, R., Abreu, I., Escudero, V., Kryvoruchko, I., Udvardi, M. K., Imperial, J., and González-Guerrero, M. (2018). *Medicago truncatula* copper transporter 1 (MtCOPT1) delivers copper for symbiotic nitrogen fixation. New Phytologist 218:696–709.

51. Sherman, Fred., Fink, G. R., Hicks, J. B., and Cold Spring Harbor Laboratory. (1983). Methods in yeast genetics. Cold Spring Harbour Lab.

52. Shin, L.-J., Lo, J.-C., and Yeh, K.-C. (2012). Copper chaperone antioxidant protein1 is essential for copper homeostasis. Plant Physiol 159:1099–1110.

53. Tejada-Jiménez, M., Castro-Rodríguez, R., Kryvoruchko, I., Mercedes Lucas, M., Udvardi, M., Imperial, J., and González-Guerrero, M. (2015). *Medicago truncatula* Natural Resistance-Associated Macrophage Protein1 is required for iron uptake by rhizobia-infected nodule cells. Plant Physiol 168:258–272.

54. Tejada-Jiménez, M., Gil-Diez, P., Leon-Mediavilla, J., Wen, J., Mysore, K. S., Imperial, J., and Gonzalez-Guerrero, M. (2017). *Medicago truncatula* Molybdate Transporter type 1 (MOT1.3) is a plasma membrane molybdenum transporter required for nitrogenase activity in root nodules under molybdenum deficiency. New Phytol 216:1223–1235.

55. Udvardi, M., and Poole, P. S. (2013). Transport and metabolism in legume-rhizobia symbioses. Annu Rev Plant Biol 64:781–805.

56. Vasse, J., de Billy, F., Camut, S., and Truchet, G. (1990). Correlation between ultrastructural differentiation of bacteroids and nitrogen fixation in alfalfa nodules. J. Bacteriol. 172:4295–4306.

57. Vernoud, V., Journet, E. P., and Barker, D. G. (2007). MtENOD20, a Nod factor-inducible molecular marker for root cortical cell activation. Mol Plant Microbe Interact 12:604–614.

58. Wong, P. C., Waggoner, D., Subramaniam, J. R., Tessarollo, L., Bartnikas, T. B., Culotta, V. C., Price, D. L., Rothstein, J., and Gitlin, J. D. (2000). Copper chaperone for superoxide dismutase is essential to activate mammalian Cu/Zn superoxide dismutase. Proc Nat Acad Sci USA 97:2886–2891.

59. Wood, W. B. (1966). Host specificity of DNA produced by Escherichia coli: bacterial mutations affecting the restriction and modification of DNA. J Mol Biol 16:118– 133.

60. Xiao, T. T., Schilderink, S., Moling, S., Deinum, E. E., Kondorosi, E., Franssen, H., Kulikova, O., Niebel, A., and Bisseling, T. (2014). Fate map of *Medicago truncatula* root nodules. Development 141:3517–3528.

61. Yatsunyk, L. A., and Rosenzweig, A. C. (2007). Cu(I) binding and transfer by the N terminus of the Wilson disease protein. J Biol Chem 282:8622–8631.

