## Supplementary Materials and Methods for "Nodule-specific Cu^+^-chaperone NCC1 is required for symbiotic nitrogen fixation in *Medicago truncatula* root nodules"

### Bioinformatics

To identify candidate $Cu^{+}$-chaperones in *A. thaliana* and *M. truncatula*, BLASTP search was performed in the *M. truncatula* Genome Project site (http://www.jcvi.org/medicago/index.php). Homologous protein sequences of these organisms were obtained from Uniprot database (http://www.uniprot.org/blast), National Center for Biotechnology Information (NCBI) (https://blast.ncbi.nlm.nih.gov/Blast.cgi?PAGE=Proteins) and from The Arabidopsis Information Resource (TAIR) (https://www.arabidopsis.org). Protein sequence comparison was carried out using ClustalW (http://www.ebi.ac.uk/Tools/msa/clustalw2/). Unrooted tree building and visualization were performed with MEGA7 (using the Neightbour Joining algorithm) and FigTree (http://tree.bio.ed.ac.uk/software/figtree/), respectively. The accessions used for *A. thaliana* were At1g12520, At1g66240, At3g56240, At5g26690, At5g60800, At1g29000, At2g36950, At5g03380, At5g63530, At3g02960, At5g24580, At1g71050, At5g17450, at1g22990, At4g39700, At4g08570, At4g35060, At4g38580, At5g66110, At3g06130, At5g19090, At3g05220, At1g56210, At5g27690, At1g23000, At1g12520, At3g05920, At4g10465, At3g56891, At5g02600, At2g37390, At3g53530, At1g51090, At1g06330, At1g29100, At2g18196, At2g28660, At3g13140, At3g21490, At3g24450, At4g15562, At4g23882, At5g05365, At5g37860, At5g50740; for *M. truncatula* the accessiones were Medtr3g111350, Medtr4g101820, Medtr7g013660, Medtr1g092670, Medtr4g133900, Medtr5g097550, Medtr1g078090, Medtr4g094280, Medtr4g062450, Medtr6g086020, Medtr6g086020, Medtr7g108560, Medtr1g082650, Medtr1g083310, Medtr7g100450, Medtr1g067560, Medtr7g113110, Medtr2g095480, Medtr4g057765, Medtr8g010540, Medtr5g069180, Medtr2g026685, Medtr3g099040, Medtr3g099030, Medtr8g096700, Medtr3g435930, Medtr1g107565, Medtr1g107565, Medtr1g063210, Medtr0041s0140, Medtr7g101930, Medtr2g436830, Medtr4g119820, Medtr4g117220, Medtr4g117140, Medtr2g022240, Medtr2g022250, Medtr5g020960, Medtr5g055020, Medtr3g117890, Medtr4g073040, Medtr5g022620, Medtr3g073670, Medtr3g087770, Medtr8g091420, Medtr5g025150, Medtr1g030630, Medtr7g083060, Medtr6g003990, Medtr6g026890, Medtr2g091235, Medtr4g120750.

**$MtNCC1_{1\text{-}78}$ $Cu^+$ loading**

$Cu^+$ loading of $MtNCC_{1\text{-}78}$ was carried out by incubating the protein in the presence of a 10-fold molar excess of $CuSO_4$, 100 mM Tris (pH 8.0), 150 mM NaCl, and 10 mM freshly prepared ascorbate for 5 min at room temperature with gentle agitation. The unbound $Cu^+$ was removed by passing through a Sephadex G-10 column (Sigma). The amount of $Cu^+$ bound was quantified using an Atomic Absorption Spectrophotometer (AAS) (ContrAA 800 spectrometer, Analytik Jena), and relativized to protein concentration.
