## Supplementary material for "Nodule-specific Cu^+^-chaperone NCC1 is required for symbiotic nitrogen fixation in *Medicago truncatula* root nodules": Fig S1

**Figure S1**

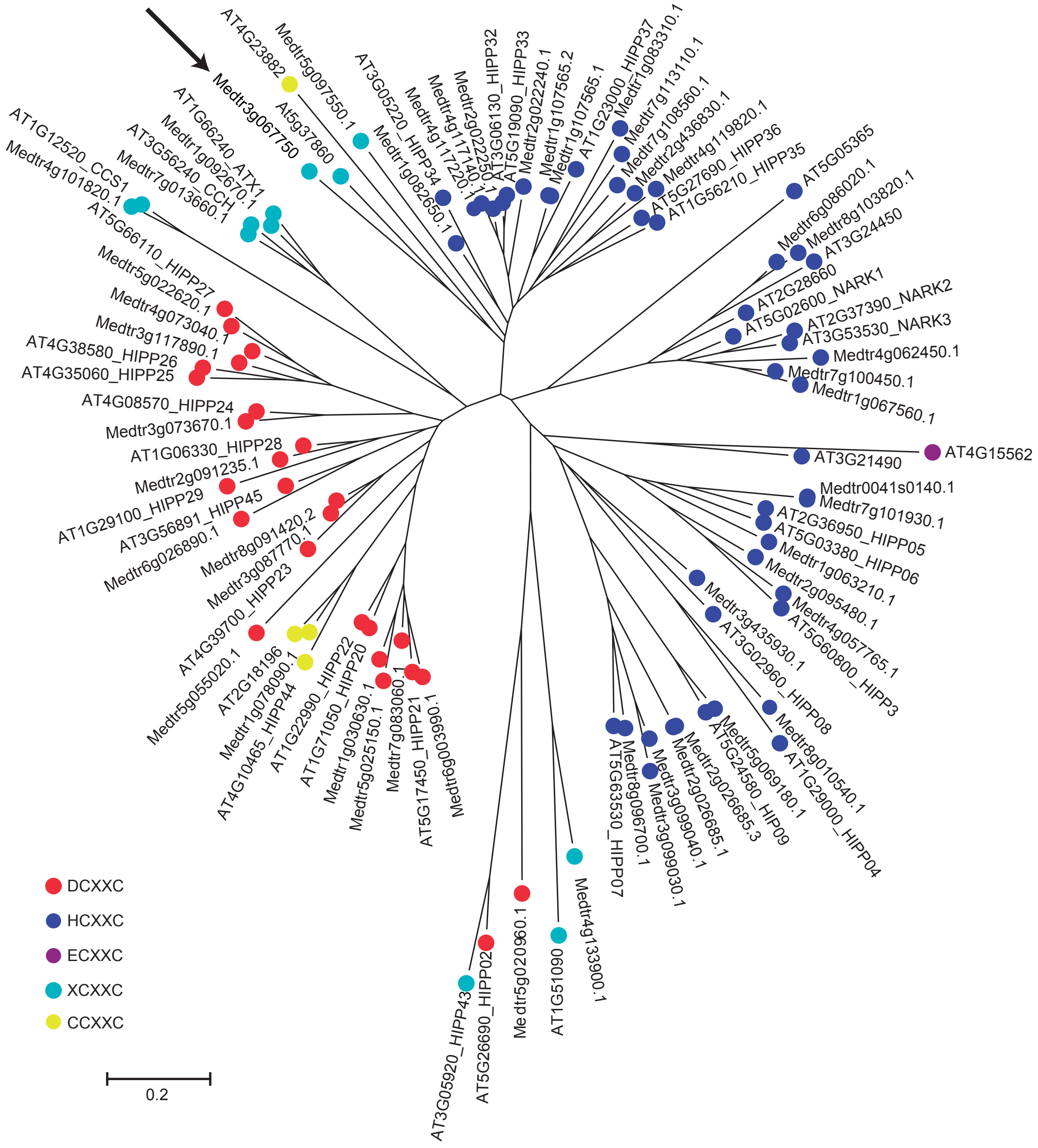


**Figure S1. Putative Cu$^{+}$-chaperones in *A. thaliana* and *M. truncatula*.** The different colours indicate the different variations of the Cu+-binding CXXC motif found in these proteins.
