## Supplementary material for "Nodule-specific Cu^+^-chaperone NCC1 is required for symbiotic nitrogen fixation in *Medicago truncatula* root nodules": Fig S2

**Figure S2**

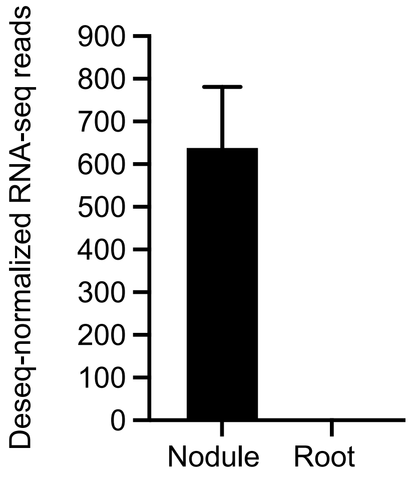


**Figure S2. *MtNCC1* expression in nodules and roots.** Data was collected from the Symbimics database (https://ia.nt.toulouse.inra.fr/symbimics/).
