## Supplementary material for "Nodule-specific Cu^+^-chaperone NCC1 is required for symbiotic nitrogen fixation in *Medicago truncatula* root nodules": Fig S3

**Figure S3**

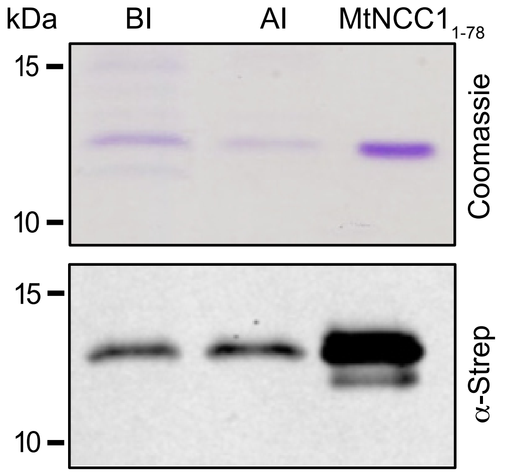


**Figure S3. Purification of MtNCC1$_{1\text{-}78}$.** Immunodetection of Twin-Strep-tagged MtNCC1$_{1\text{-}78}$ in crude *E. coli* Bl21 protein extracts before the induction (BI) and after the induction (AI) and after purification (lower panel). Higher panel corresponds to the Coomassie Brilliant Blue staining of these protein extracts.
