## Supplementary material for "Nodule-specific Cu^+^-chaperone NCC1 is required for symbiotic nitrogen fixation in *Medicago truncatula* root nodules": Fig S4

**Figure S4**

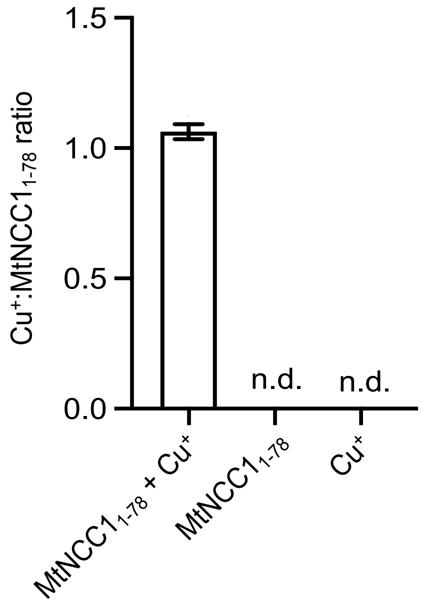


**Figure S4. $Cu^+$-binding to MtNCC1$_{-78}$.** Determination of copper content in desalted fractions of MtNCC1$_{1\text{-}78}$ previously incubated with 10 x molar excess of $Cu^+$, of MtNCC1$_{1\text{-}78}$ that has not been incubated with $Cu^+$, and a control with only the 10 x molar excess of $Cu^+$ (control for desalting column). Data are the mean ± SE (n = 3). n.d. stands for not-detected.
