## Supplementary material for "Nodule-specific Cu^+^-chaperone NCC1 is required for symbiotic nitrogen fixation in *Medicago truncatula* root nodules": Fig S5

### Figure S5

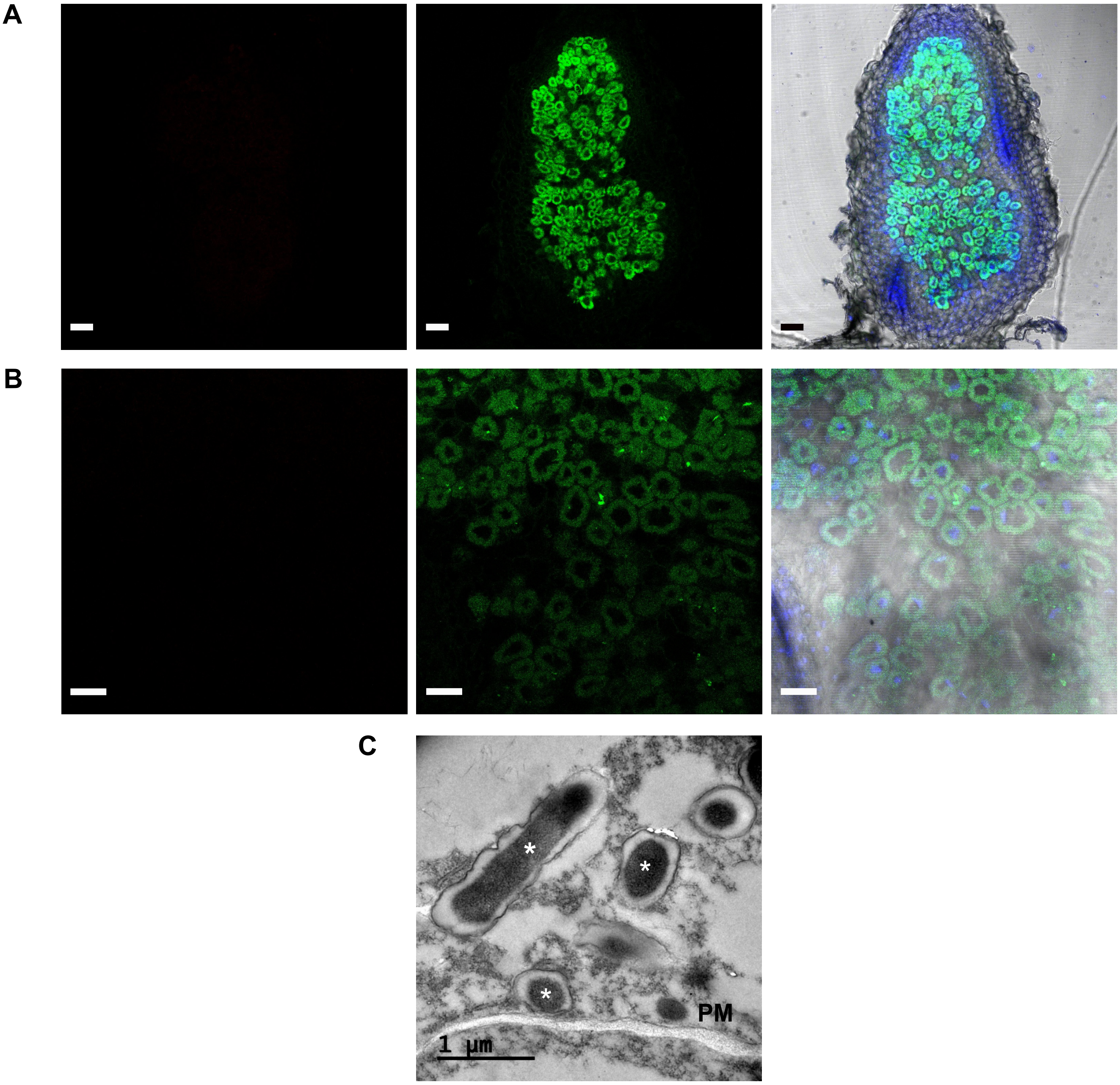


**Figure S5. Control for immunolocalization.** A, Confocal image of wild type 28 dpi *M. truncatula* nodule sections incubated Alexa 594-conjugated antibody (red, when positive, left panel), on nodules inoculated with GFP-expressing *S. meliloti* (green, central panel), and stained with DAPI (blue). Right panel is the overlaid image of the transillumination image with the DAPI, Alexa 594, and GFP channels. Bars = 100 µm. B, Detailed view of the differentiation zone. Bar = 60 µm. C, Wild type 28 dpi *M. truncatula* nodule sections incubated with an anti-HA gold-conjugated antibody. * indicates bacteroids. Bar = 1 µm.
