## Supplementary material for "Nodule-specific Cu^+^-chaperone NCC1 is required for symbiotic nitrogen fixation in *Medicago truncatula* root nodules": Fig S6

**Figure S6**

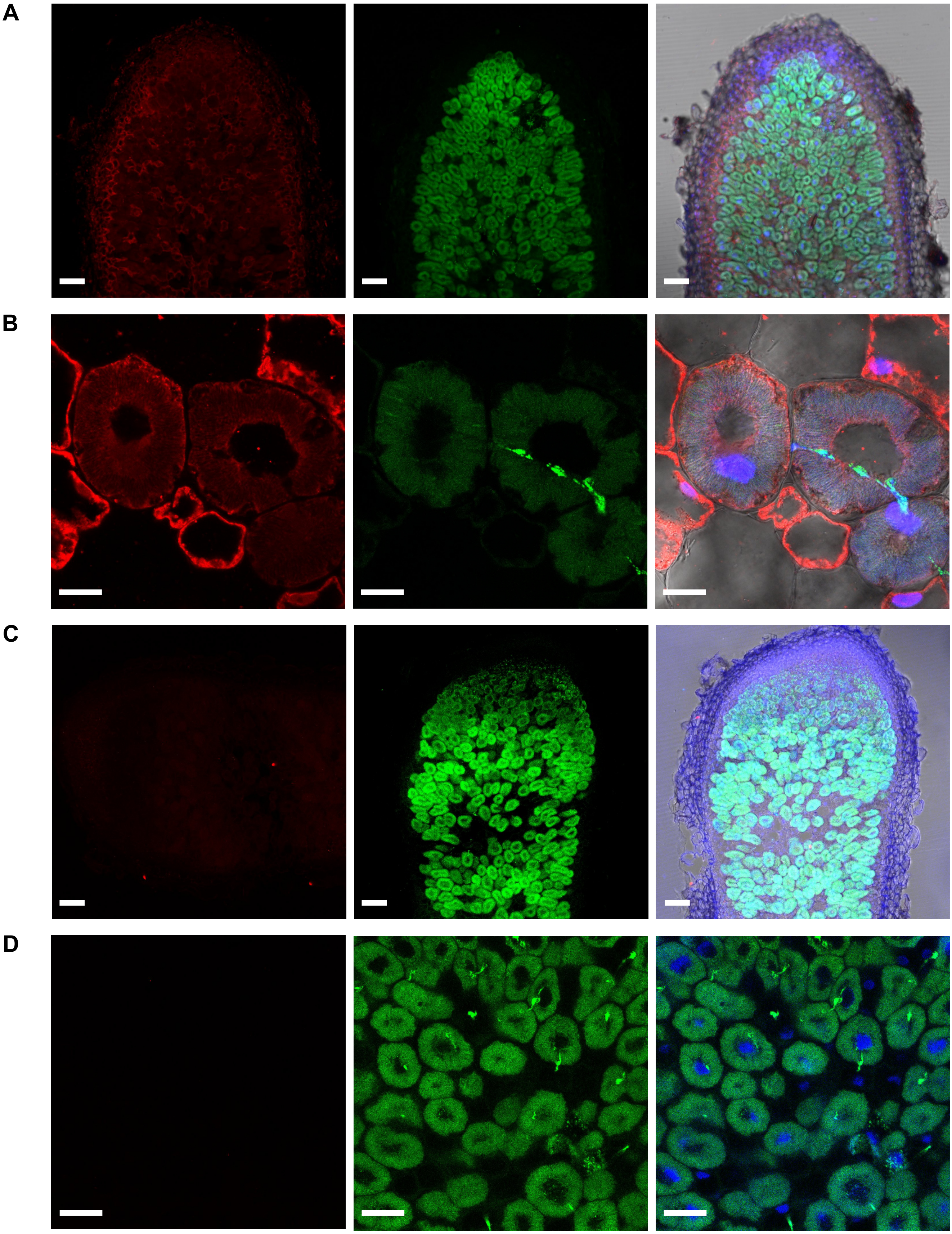


**Figure S6. Immunolocalization of MtNCC1$_{1\text{-}78}$-HA.** A, Longitudinal section of 28 dpi *M. truncula* nodules expressing MtNCC1$_{1\text{-}78}$ fused to three HA domains and regulated by its own promoter. HA-tagged proteins were detected with using an Alexa594 conjugated antibody (red, left panel). Nodules were colonized by a GFP-expressing *S. meliloti* (green, central panel). DNA was stained with DAPI (blue) and overlaid with the previous two channels and the transillumination signal (right panel). Bars = 100 µm. B, Closer view of rhizobia-infected cell in zone II. Bars = 20 µm. C, D, negative controls for MtNCC1$_{1\text{-}78}$ immunolocalization. Bars = 100 and 20 µm, respectively
