## Supplementary material for "Nodule-specific Cu^+^-chaperone NCC1 is required for symbiotic nitrogen fixation in *Medicago truncatula* root nodules": Fig S7

**Figure S7**

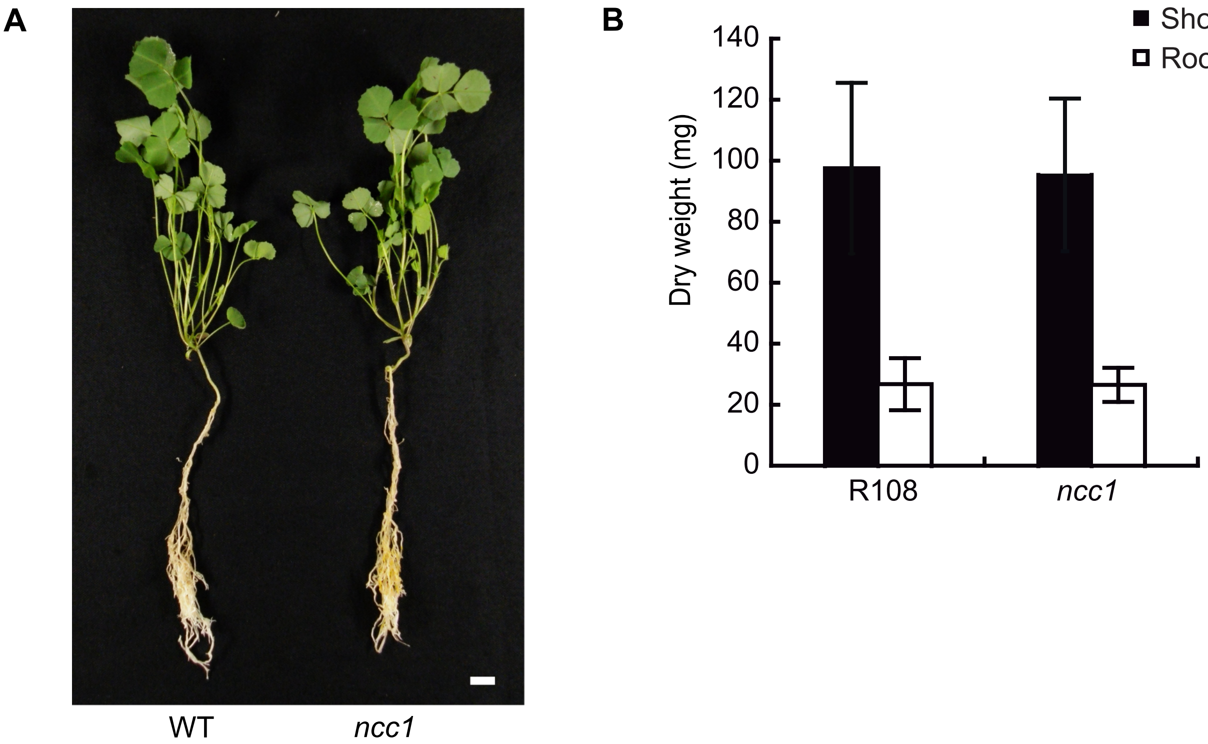


**Figure S7. MtNCC1 is not necessary during non-symbiotic conditions.** A, Representative images of wild type (WT) and *ncc1* mutant non-inoculated *M. truncatula* plants irrigated every two weeks with 2 mM ammonium nitrate. Bar = 1 cm. B, Dry weight of shoots and roots from WT and *ncc1* mutant non-inoculated M. truncatula plants irrigated every two weeks with 2 mM ammonium nitrate. Data are the mean ± SE (n = 10).
