## Supplementary figures and images for "Nodule-specific Cu^+^-chaperone NCC1 is required for symbiotic nitrogen fixation in *Medicago truncatula* root nodules"

### Fig S8

#### Figure S8

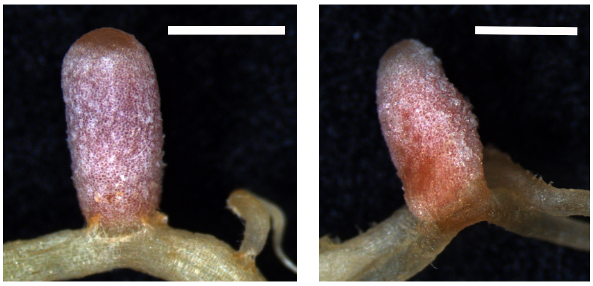

WT *ncc1*

**Figure S8. Representative images of 28 dpi wild type (WT) and *ncc1* mutant nodules.** Bar = 1 mm.
