## Supplementary material for "Nodule-specific Cu^+^-chaperone NCC1 is required for symbiotic nitrogen fixation in *Medicago truncatula* root nodules": Fig S9

**Figure S9**

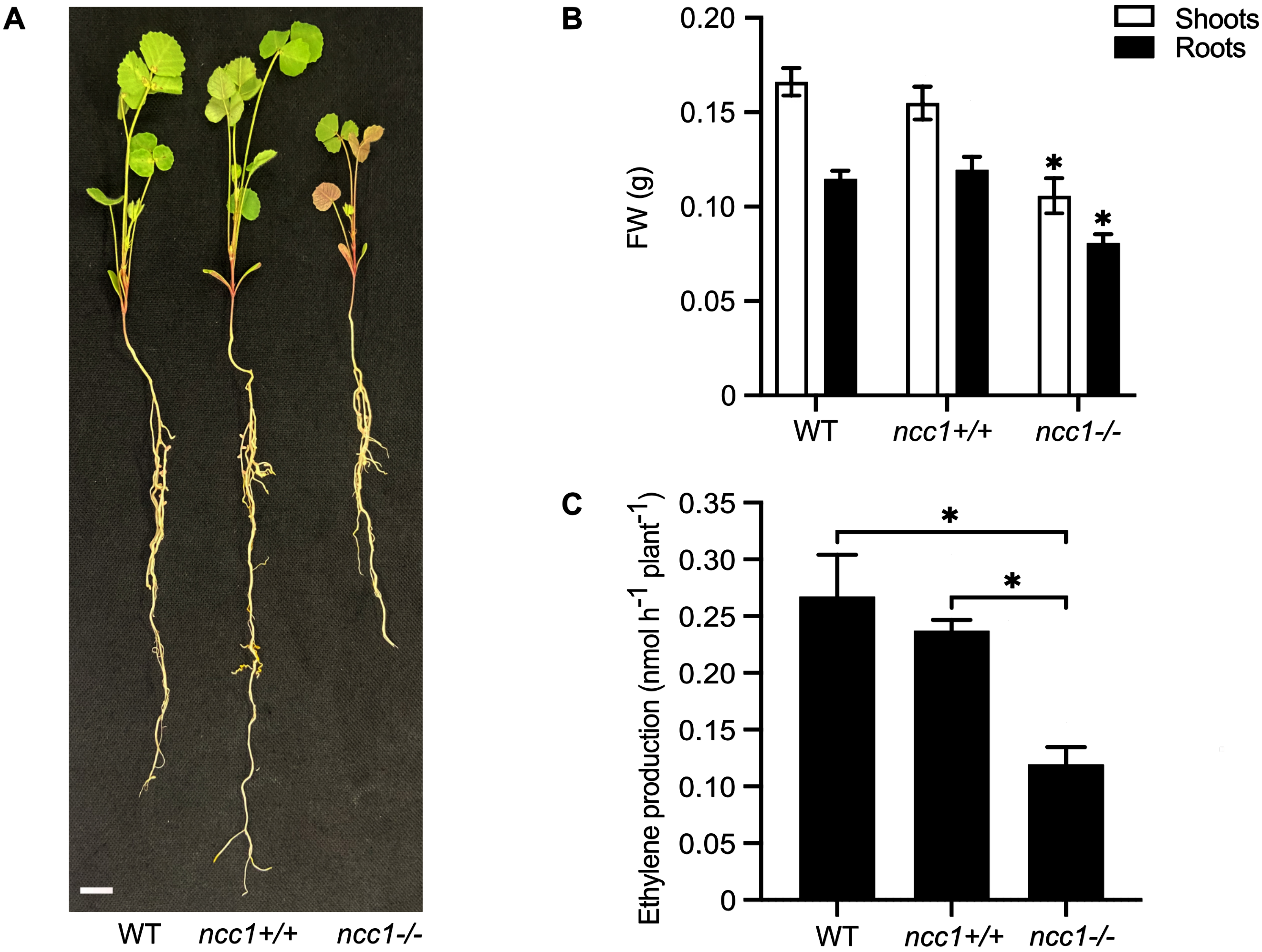


**Figure S9. Growth of 28 dpi wild type (WT), and wild type (+/+) and mutant (-/-) *ncc1* segregants.** A, Representative images of 28 dpi WT, *ncc1*+/+, and *ncc1*-/- plants. Bars = 1 cm. B, Fresh weight (FW) of 28 dpi WT, *ncc1*+/+, and *ncc1*-/- plants. Bars represent the mean ± SE (n = 35). C, Nitrogenase activity of 28 dpi WT, *ncc1*+/+, and *ncc1*-/- plants. Bars represent the mean ± SE of three sets of 5 pooled plants each.
