## Supplementary material for "Nodule-specific Cu^+^-chaperone NCC1 is required for symbiotic nitrogen fixation in *Medicago truncatula* root nodules": Fig S10

### Figure S10

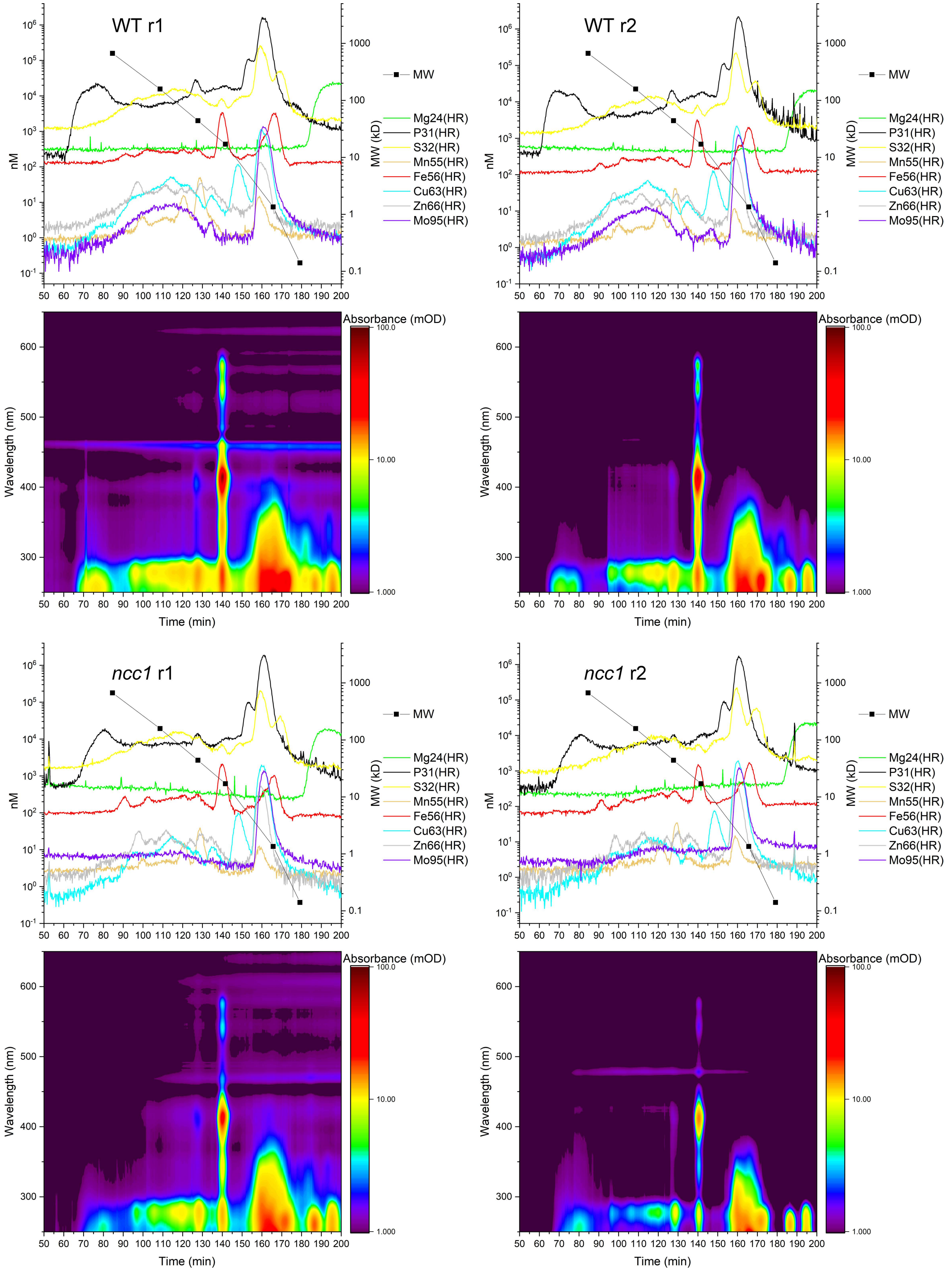


**Figure S10. Complete HPLC-ICPsfMS chromatograms.** These graphs show the complete time range of the samples from which Figure 6 was constructed by focusing on the MW range where the largest differences were observed between the WT and the *ncc1* mutant.
