## Supplementary material for "Nodule-specific Cu^+^-chaperone NCC1 is required for symbiotic nitrogen fixation in *Medicago truncatula* root nodules": Fig S11

**Figure S11**

A

B

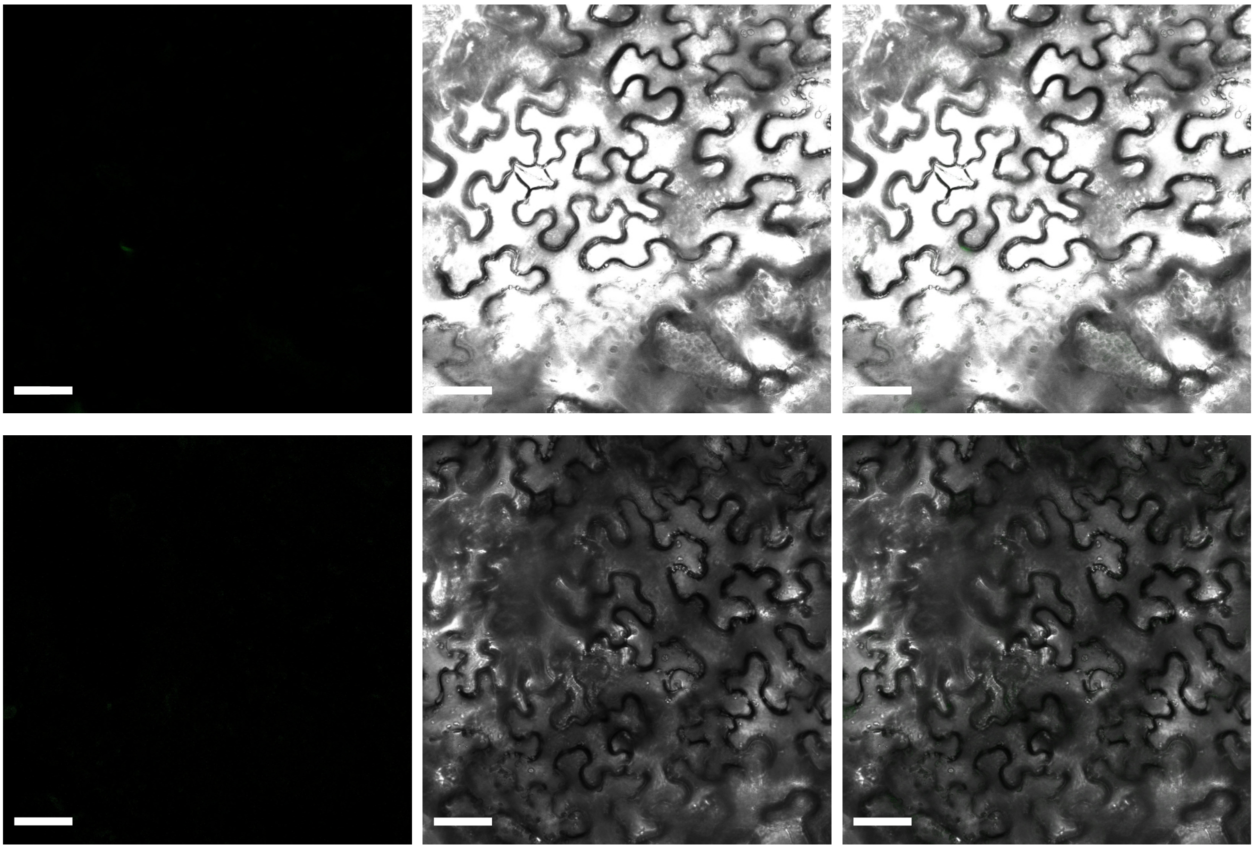

**Figure S11. Negative controls for BiFC**. A, Co-agroinfiltration of $MtNCC1_{1\text{-}78}$ in pNXGW with empty pCXGW. B, Co-agroinfiltration of $MtNCC1_{1\text{-}78}$ in pNXGW with empty pXCGW. Left panels show where the GFP signal should appear in case of interaction; middle panels are the transillumination, and right panels correspond to the overlay of the two channels. Bars = 50 μm
