## Supplementary material for "Nodule-specific Cu^+^-chaperone NCC1 is required for symbiotic nitrogen fixation in *Medicago truncatula* root nodules": Table S5

**Supplemental Table 5:** Primers used in this study.

| **Primer** | **Sequence** | **Use** |
| --- | --- | --- |
| 5 MtUb v4qF | ATTCTTCACATGCGGCGATTTAC | qRT-PCR of *ubiquitin carboxylterminal*  *hydrolase* |
| 3 MtUb v4qR | TTTCTCATTTGCTTTTGGTGTTGG | qRT-PCR of *ubiquitin carboxylterminal*  *hydrolase* |
| 5 MtNCC q1191 F | AGATGATGATGATGATGATGATGA | qRT-PCR of *MtNCC1* *(Medtr3g067750)* |
| 3 MtNCC q1279 R | CCTTTGTCTTATGAAACTTTTGAC | qRT-PCR of *MtNCC1 (Medtr3g067750)* |
| 5 MtCCS1 ATG  pDR196 tail F | CACATTCAAAAGAAAGAAAAAAAATATACCCCAGCCTCGAATGTCAACAAATGAACATGAGT | *MtNCC11/ MtNCC_1-78_* expression in yeast |
| 3 MtCCS1 585  pDR196 tail R | GACTTGACCAAACCTCTGGCGAAGAAGTCCAAAGCTGGATTTAATTTGAAGGGTGATAACCAT | *MtNCC1* expression in yeast |
| 3 MtCCS1 237  pDR196 tail R | GACTTGACCAAACCTCTGGCGAAGAAGTCCAAAGCTGGATTTCATTCAAAATCTCTGCATGTTT | *MtNCC1_1-78_* expression in yeast |
| 5 MtNCCp -2177  GW F | GGGGACAAGTTTGTACAAAAAAGCAGGCTGCCAAAGTAATTGAATGTCTAGT | Cloning of *MtNCC1* promoter in pGWB3/ promoter + coding sequence in pGWB13 |
| 3 MtNCCp -1 GW R | GGGGACCACTTTGTACAAGAAAGCTGGGTAGGCTTGAAGCTAGAATATAAGAAAAG | Cloning of *MtNCC1* promoter in pGWB3 |
| 3 MtNCC +1498 GW R | GGGGACCACTTTGTACAAGAAAGCTGGGTAATTTGAAGGGTGATAACCATG | Cloning of *MtNCC1* promoter + coding sequence in pGWB13 |
| 5 NCC N-pET16b tail F | AAAAAGAAAATCTTTATTTTCAAGGTCATATGATGTCAACAAATGAACATGAGTC | Cloning of *MtNCC1_1-78_* in modified pET16b |
| 3 NCC trun  N-pET16b tail R | GGGCTTTGTTAGCAGCCGGATCCTTATTCATTCAAAATCTCTGCATGTTTAC | Cloning of *MtNCC1_1-78_* in modified pET16b |
| MtNCC FW GW | GGGGACAAGTTTGTACAAAAAAGCAGGCTTCATGTCAACAAATGAACATGAG | Cloning of *MtNCC1_1_*_-78_ in pNXGW |
| MtNCC_1-78_ STOP RV GW | GGGGACCACTTTGTACAAGAAAGCTGGGTCTTATTCATTCAAAATCTCTGCATG | Cloning of *MtNCC1_1_*_-78_ in pNXGW |
| Medtr7g105830  FW GW | GGGGACAAGTTTGTACAAAAAAGCAGGCTTCATGGCCTGCTCAGCTCCATCT | Cloning of *Medtr7g105830* in pCXGW/pXCGW |
| Medtr7g105830 STOP RV GW | GGGGACCACTTTGTACAAGAAAGCTGGGTCCTACACTGCAGAAAAGTACTCTTT | Cloning of *Medtr7g105830*  in pCXGW |
| Medtr7g105830  RV GW | GGGGACCACTTTGTACAAGAAAGCTGGGTCCACTGCAGAAAAGTACTCTTT | Cloning of *Medtr7g105830*  in pXCGW |
| Medtr2g046710  FW GW | GGGGACAAGTTTGTACAAAAAAGCAGGCTTCATGGCAACAGAAACTTTCCTA | Cloning of *Medtr2g046710*  in pCXGW/pXCGW |
| Medtr2g046710 STOP RV GW | GGGGACCACTTTGTACAAGAAAGCTGGGTCTTAGGCAGTGGCTAACTTTCCACC | Cloning of *Medtr2g046710*  in pCXGW |
| Medtr2g046710  RV GW | GGGGACCACTTTGTACAAGAAAGCTGGGTCGGCAGTGGCTAACTTTCCACC | Cloning of *Medtr2g046710*  in pXCGW |
| Medtr1g083950  FW GW | GGGGACAAGTTTGTACAAAAAAGCAGGCTTCATGGCTGGCATAACAGAAAAC | Cloning of *Medtr1g083950*  in pCXGW/pXCGW |
| Medtr1g083950  STOP RV GW | GGGGACCACTTTGTACAAGAAAGCTGGGTCTTAATTATCTCCTCCAGTGGTTGA | Cloning of *Medtr1g083950*  in pCXGW |
| Medtr1g083950  RV GW | GGGGACCACTTTGTACAAGAAAGCTGGGTCATTATCTCCTCCAGTGGTTGA | Cloning of *Medtr1g083950*  in pXCGW |
| Medtr2g076070  FW GW | GGGGACAAGTTTGTACAAAAAAGCAGGCTTCATGTCTAACATGGTTGCAAAA | Cloning of *Medtr2g076070*  in pCXGW/pXCGW |
| Medtr2g076070  STOP RV GW | GGGGACCACTTTGTACAAGAAAGCTGGGTCTTAATTGGAGAATGGACATGCTGA | Cloning of *Medtr2g076070*  in pCXGW |
| Medtr2g076070  RV GW | GGGGACCACTTTGTACAAGAAAGCTGGGTCATTGGAGAATGGACATGCTGA | Cloning of *Medtr2g076070*  in pXCGW |
